# Multi-resolution humanness scoring of antibodies via a region-isolated language model

**DOI:** 10.64898/2026.10.23.761395

**Authors:** Huaqing Liu, Peiyi Chen, Shuxian Zhou, Bin Wei, Shuwen Deng, Lixiong Zhang, Xinlin Liu, Shun Zhou, Lanqing Han, Li Ye

## Abstract

Antibodies recognized as foreign can elicit anti-drug antibodies that compromise efficacy and safety, making humanness scoring, estimating how human-like a sequence appears, an early, repeated step in antibody development. Widely used scores reduce a variable domain to a single framework-dominated number, leaving the complementarity-determining regions (CDRs), grafted during humanization and diversified by generative models, weakly supervised and rarely audited. Here we introduce IgScope, a 23-million-parameter antibody language model trained from scratch that scores humanness at chain, IMGT-region, and residue resolution; a region-isolated attention module confines attention to within-region pairs, and chimeric-sequence training supplies region-level supervision. On 2.38 million held-out chains, masking all framework tokens leaves CDR accuracy unchanged, and back-mutated murine residues in human frameworks are detected at 93.4% accuracy. At the chain level, IgScope matches the strongest published correlation with clinical anti-drug-antibody incidence (r = ***−*0.567**, n = 217) and outperforms a same-protocol recomputation. Region-level scoring exposes what chain-level scoring conceals: antibodies designed de novo by four generative models pass whole-chain humanness (0.966–0.988), while their CDRs score far below natural (P ***<*** 1e-30) in every chain class. Quality control turns the same resolution to the mirror-image question: in humanized-mouse platforms, whose domains are human by design, IgScope scores the domain as human and, in HUGO-Nano chimeric VHHs, isolates only the FR2 signal of four camelid hallmark residues deliberately retained to stabilize the VHH. IgScope gives antibody humanization a verified foundation and generative design an audit.

## Introduction

Humanness assessment is a routine gate in therapeutic antibody development. Antibodies of animal origin, and increasingly antibodies designed by computational generators, are scored for how closely their sequences resemble natural human repertoires before they enter the long path of candidate optimization. Scores in common use include OASis, which searches the Observed Antibody Space for similar human sequences ^1,2^; T20, which averages a chain’s identity to its 20 closest human matches in a reference database ^3^; AbNatiV, a set of chain-type-specific variational-autoencoder nativeness models (one each for heavy, kappa and lambda chains) ^4^; and HumAb, a V-gene-family random-forest classifier ^5^. Each returns one number per chain. That number is dominated by the frameworks, which constitute roughly three-quarters of a variable domain and are strongly conserved within a species, whereas the CDRs— grafted during humanization, diversified by generative models and retained from the donor species in chimeric constructs—enter only through an average. A chain can therefore receive a reassuring score while the region most likely to be foreign has not been examined in its own right.

Two developments make this compression consequential. First, generative models now design antibodies and nanobodies against user-specified epitopes ^6–9^. In the common workflow the framework is supplied by the user—typically as a human germline—and the CDRs are generated de novo, so the designed CDR is the principal new sequence in the molecule. Second, a principal route to anti-drug antibodies is T-cell help: CD4^+^ T cells recognize short linear peptides displayed by MHC class II ^10–12^. T-cell recognition is therefore local—it reads peptides, not intact chains—so a single non-human stretch can create an epitope even when the rest of the variable domain is human. In conventional antibodies the non-human segments are inherited from the donor and are the object of humanization; in designed antibodies they are the newly generated CDRs. A chain-level average cannot report whether such a segment exists. The scores above were not built to answer this question. They are trained and calibrated on natural chains whose labels are chain-level, and in such data the framework is both abundant and strongly species-discriminative: species-specific positions in the framework carry a signal that is easy to learn. Under these conditions a learned model can reach high accuracy from framework markers alone without representing the species features of CDRs—a classic shortcut-learning failure. For alignment- and identity-based scores the imbalance is structural rather than learned, because three-quarters of the positions they average over lie in the framework. Protein language models inherit the same bias: AntiBERTy, which carries species- and chain-type-prediction heads trained on natural repertoires, predicts both at the chain level ^13^. A model can therefore be accurate and still have learned little about the region that matters, a form of laziness that no chain-level benchmark exposes. Three questions follow.

Is the species signal confined to the framework, or do the CDRs carry it as well? Do existing whole-chain scorers use that CDR information, or do they reach their accuracy from the framework alone? And can humanness be reported at the resolution at which engineering decisions are made, the region and the residue? None of the three has, to our knowledge, been tested directly.

We answer all three with IgScope, a 23-million-parameter single-chain antibody language model trained from scratch on 119 million natural heavy, kappa, lambda and VHH chains, which outputs the probability of human origin at three resolutions (chain, the seven IMGT regions FR1–CDR3–FR4, and residue) in one forward pass (Fig. 1). The central difficulty is not model capacity but shortcut learning: because frameworks carry a strong species signal, a naively trained model has every incentive to classify a CDR from its framework context. IgScope removes that option architecturally. A region-isolated Transformer module restricts attention to within-region tokens, so the representation of a CDR, and hence its humanness score, cannot depend on framework tokens. Chimeric-sequence training then supplies the supervision that natural sequences lack: synthetic cross-species grafts make up 30% of the training stream, receive region- and residue-level labels from the donor species, and are excluded from the chain-level loss. Two evolution-aware auxiliary tasks (ancestral germline prediction and mutation-process prediction) add dense germline-alignment supervision to the residue head.

**Fig. 1.**
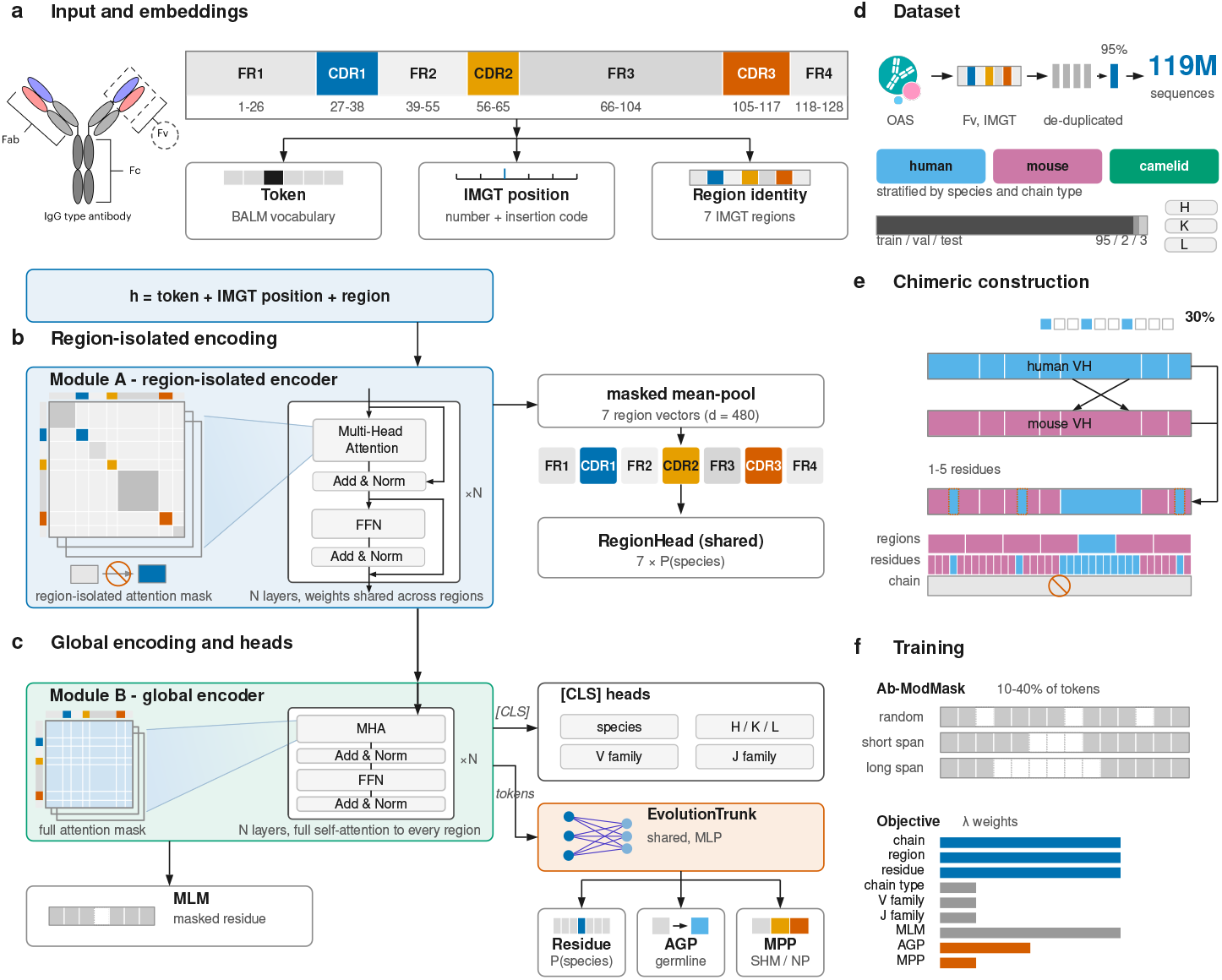
IgScope: region-isolated encoding for multi-resolution species-origin scoring. (a) One unpaired antibody variable chain (VH, VL or VHH) in IMGT numbering, with the seven IMGT regions. Residue identity, IMGT position and region identity are summed into the input embedding; chain type, V/J gene and species are prediction targets, never inputs. (b) Module A: a same-region attention mask (crossed arrow) prevents framework (FR) context from reaching complementarity-determining region (CDR) representations; each region is mean-pooled and scored by a shared region head. (c) Module B: full attention; its class token feeds the chain-level heads, and its token states feed the residue, germline-reconstruction (AGP) and mutation-process (MPP) heads through a shared trunk. (d) Training corpus: human, mouse and camelid repertoires, 119M chains. (e) Chimeric training: regions from two species, with the chain-level label withheld. (f) Ab-ModMask corruption and the nine-term training objective.

Across 2.38 million held-out natural chains, IgScope’s region scores are verifiably local. Masking every framework token leaves CDR classification accuracy unchanged (drop = 0.0000), and the mirror ablation degrades AntiBERTy’s species call far more than masking its CDRs (AUC-ROC 0.672 versus 0.873), which locates that model’s species signal in the framework. Murine residues reintroduced into human frameworks are identified at 0.934 accuracy. At chain level, IgScope matches the strongest published correlation with clinical anti-drug-antibody incidence (Pearson r = *−*0.567, n = 217), exceeds every other published baseline on that benchmark, and classifies anti-drug-antibody status better than four HLA class II epitope-based predictors on the same antibodies (ROC-AUC 0.807); residue-level features describing non-human patches correlate with incidence as strongly as whole-chain scores, consistent with immunogenicity arising from local segments rather than global dissimilarity. The resolution matters where it is applied. Antibodies designed de novo by four generative models pass whole-chain humanness (0.966–0.988) while their CDRs score far below natural in heavy, light and VHH chains alike (P *<* 1e-30). The mirror-image question of quality control is answered by the same scores: humanized-mouse platforms engineered to carry human variable domains score as human across the domain, and in chimeric VHHs IgScope isolates only the FR2 signal of the four camelid hallmark residues deliberately retained on an otherwise human framework, which chain-level scores cannot see. These scores are computational hypotheses about immunogenic potential and require experimental confirmation; within that boundary, IgScope gives generative antibody design an audit and humanization a foundation that can be checked at the region and residue level.

## Results

### A region-isolated language model scores humanness at three resolutions

IgScope is a 23-million-parameter Transformer encoder trained from scratch on 119 million natural antibody chains (Fig. 1; Methods). A single chain is embedded as the sum of residue-token, factorized IMGT-position and region embeddings; species, chain type and V/J gene are never given as inputs and serve only as prediction targets. The first encoder module (Module A) applies a same-region attention mask (every token attends only to tokens of its own IMGT region, never across regions), and its per-region representations are scored by a parameter-shared region head. The second module (Module B) attends over the full sequence; its class token drives the chain-level species head and auxiliary heads, and its per-token states feed a shared evolution trunk from which the residue, germline-reconstruction (AGP) and mutation-process (MPP) heads are computed (Methods).

Internal validation places the heads near their task ceilings. On 2,379,517 natural chains held out from training, the chain-level species head reaches three-way accuracy 0.9998 (macro-F1 0.9997), the region head reaches 0.9618–0.9771 accuracy on CDR1/2/3, and the auxiliary heads reach AUPRC 0.999997 at the residue level, 0.9925 token accuracy for ancestral germline reconstruction and 0.9927 three-class accuracy for mutation-process prediction (Extended Data Table 1). High accuracy is not, however, evidence of locality: a chain-level model that reads species information from the framework would report the same chain-level accuracy, and a region score can be accurate while being computed from context rather than from the region’s own sequence.

We therefore asked whether the fine-grained scores are *local* —whether a region score is computed from that region’s own sequence rather than from its framework context. Replacing every framework token with a mask token and re-scoring leaves CDR accuracy unchanged (drop = 0.0000 for CDR1/2/3; Fig. 2a): to measurement precision, each region score is a function of its own region’s sequence. On synthetic back-mutated chains (murine residues reintroduced into human frameworks), residue-level identification accuracy is 0.934, versus 0.997 on unmutated vernier positions ^14^ (Fig. 2b). This sensitivity is a direct consequence of the training design: on chimeric constructs, isolated donor-species residues must be flagged at exactly their positions, so the residue head learns to mark a murine residue inside a human framework (Fig. 2b).

**Fig. 2.**
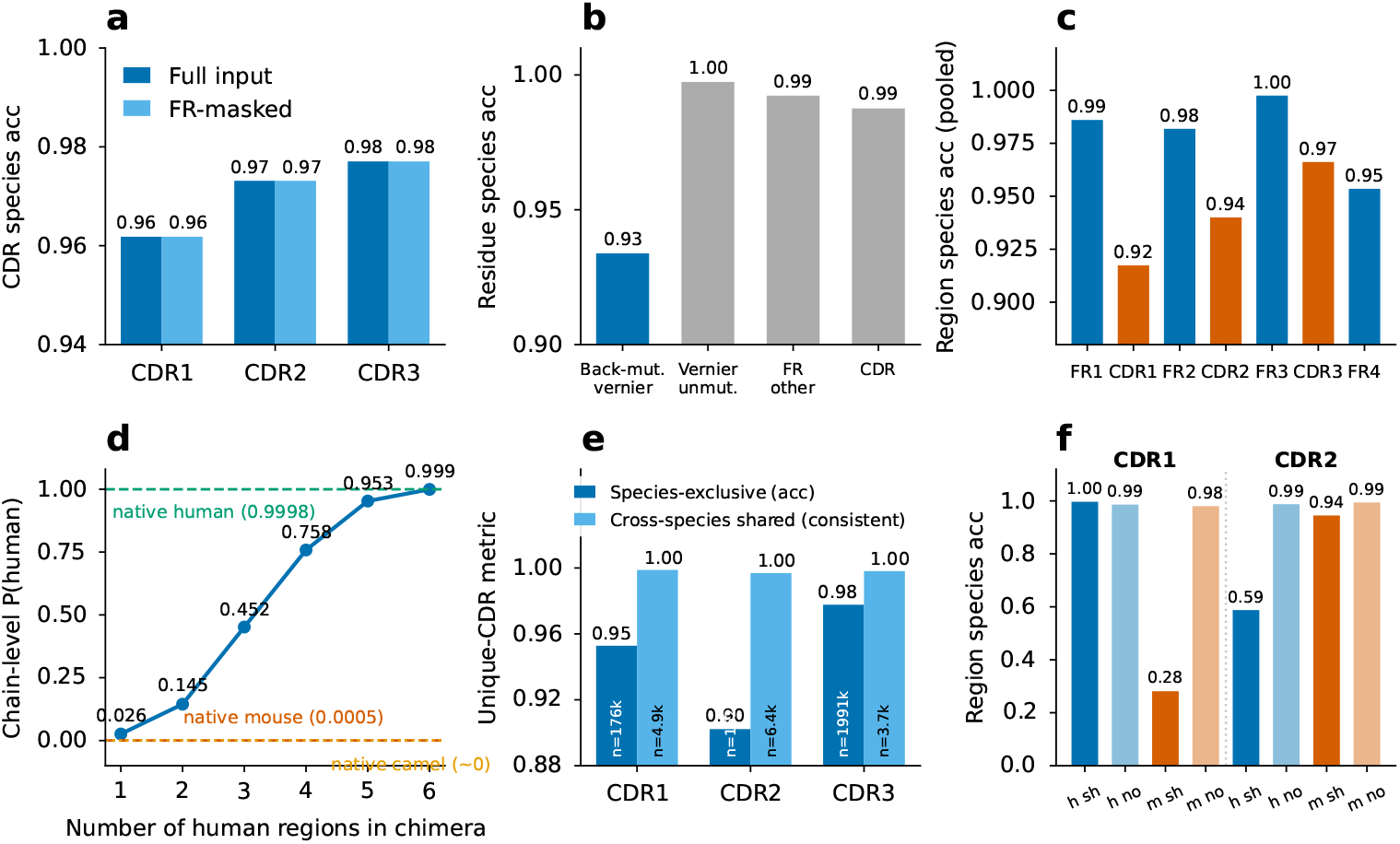
Multi-resolution diagnostic suite. a, Framework-masked CDR test: replacing all framework tokens with mask tokens leaves CDR accuracy unchanged. b, Back-mutation diagnostic: residue-level identification of murine residues reintroduced into human frameworks, with unmutated vernier, other framework and CDR positions as controls. c, Region accuracy on synthetic chimeras. d, Chain-level P(human) versus the number of human regions in chimeras, which never enter the chain loss. e, Accuracy on species-exclusive versus cross-species shared CDRs. f, Error enrichment on shared germline motifs (OR, error-enrichment odds ratio, equation (5)); OR = 131.7 for mouse CDR1 motifs read as human and 220.3 for human CDR2 motifs read as mouse; h, human; m, mouse; sh, shared; no, non-shared.

On 6,000 synthetic chimeras, pooled seven-region accuracy is 0.963 (FR-mean 0.980, CDR-mean 0.941; Fig. 2c). Although chimeras never contribute to the chain-level loss, chain-level P(human) propagates monotonically with the number of human regions (0.026 *→* 0.999 for one to six human regions; Fig. 2d), an out-of-distribution generalization consistent with additive region evidence. The residual region-level error is concentrated where error is unavoidable: on CDR1 motifs identical between human and mouse germline families, mouse accuracy falls to 0.281 (odds ratio 131.7 versus species-exclusive motifs; Fig. 2e, f), yet such a shared motif receives a single, stable species call, which falls within the true species set for more than 99.6% of shared motifs—the ceiling for any region-local classifier that sees the same sequence twice (Extended Data Fig. 1).

#### IgScope surpasses published humanness scorers on fully external repertoires, with anti-shortcut behavior preserved

We first benchmarked IgScope against published scorers under a controlled, same-protocol comparison on data none of the methods had seen. We scored a stratified subsample of the external natural set (10,000 chains per species from the 1,005,481-chain VDJServer external test set, which we curated from six BCR studies deposited in the VDJServer AIRR Data Commons ^15–22^ and which is disjoint from the training corpus; several scorers do not scale to the full million-chain set) with OASis, T20, Humatch and the native AntiBERTy species head alongside IgScope. IgScope separates human from non-human chains better than every comparator (area under the ROC curve 0.9997, versus 0.9927 for Humatch, 0.9300 for AntiBERTy, 0.9239 for OASis and 0.9228 for T20; area under the precision-recall curve 0.9994 versus 0.9879, 0.8579, 0.8672 and 0.8644; Fig. 3a, b). The separation pattern is itself diagnostic: after per-method min–max rescaling for display (Methods), IgScope and Humatch saturate (median human score approximately 1.0, non-human approximately 0), whereas OASis and T20 produce graded distributions that overlap substantially between species; the framework-ablation analyses below examine this dependence directly (Fig. 3c and Extended Data Fig. 2).

**Fig. 3.**
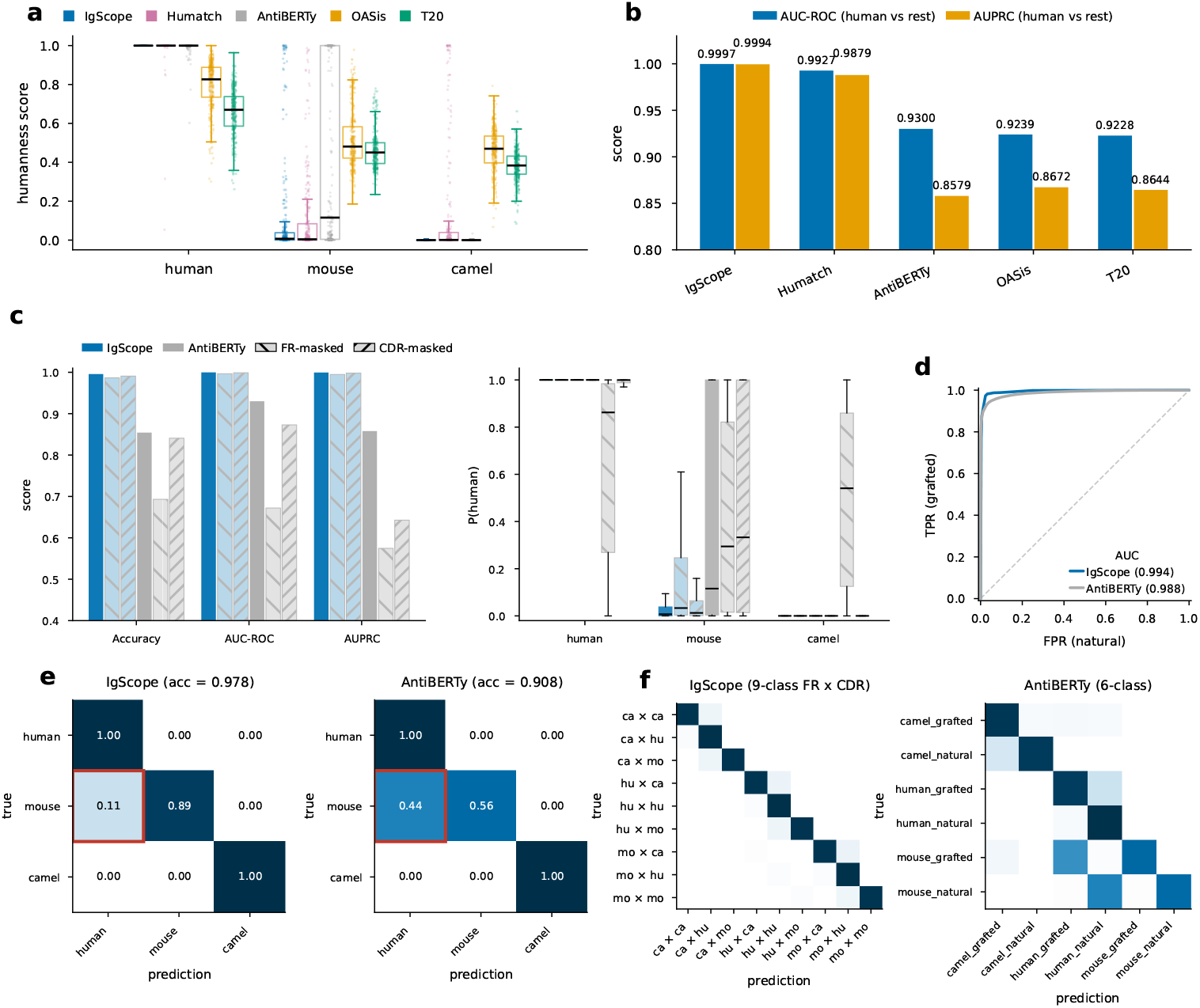
External validation: comparison with published humanness scorers and with AntiBERTy classification heads. a, b, Humanness-scorer comparison on a stratified subsample of the external natural set. a, Per-species distributions of each scorer’s native score, min–max rescaled per method to [0, 1] for display, so the axis is common but its endpoints are method-specific (native ranges in Methods); boxes, interquartile range; points, sampled chains. b, Species discrimination (AUC-ROC and AUPRC; human versus non-human). c, Mask-token ablation for the two antibody 3 language models: species classification with intact (solid), framework-masked (backslash-hatched) or CDR-masked (forward-slash-hatched) inputs, and per-species P(human) for intact, framework-masked and CDR-masked inputs (boxes, interquartile range; whiskers, 1.5 *×* range). Mask tokens cannot be ANARCI-numbered, so the all-method substitution is in Extended Data Fig. 2. d, Graft-versus-natural detection (ROC curves). e, Confusion matrices (rows normalized; red boxes, mouse-to-human cell). f, Graft composition by species class (IgScope, nine FR/CDR classes; AntiBERTy, six).

We next asked whether IgScope’s anti-shortcut properties survive this distribution shift, comparing against the native classification heads of AntiBERTy ^13^, the only antibody language model that ships species and graft heads. On the full external set, IgScope attains an accuracy of 0.9783 *±* 0.0005 (Matthews correlation 0.9657) versus AntiBERTy’s 0.9084/0.8624 (Fig. 3e). To separate CDR-based discrimination from framework reliance, we rebuilt a 30,000-chain benchmark (10,000 chains per species) and applied two complementary mask-token ablations: every framework token (FR1–FR4) replaced, or every CDR token (CDR1–CDR3) replaced. IgScope lost little under either ablation (AUC-ROC 0.9965 with frameworks masked and 0.9988 with CDRs masked, versus 0.9997 intact), whereas AntiBERTy lost most of its discrimination when frameworks were masked (AUC-ROC 0.672, AUPRC 0.575, versus 0.930 and 0.858 intact) and much of its precision–recall performance when CDRs were masked (AUPRC 0.643, AUC-ROC 0.873) (Fig. 3c)—to our knowledge the first side-by-side demonstration that an antibody species classifier’s CDR judgments survive complete framework ablation while a comparably trained language model’s degrade. The asymmetry is diagnostic: masking frameworks degrades AntiBERTy far more than masking CDRs (AUC-ROC 0.672 versus 0.873; accuracy 0.693 versus 0.841), indicating that its species call rests predominantly on framework features, whereas IgScope is essentially unchanged under either ablation and so does not depend on either region alone. Per-species medians are unmoved for IgScope (human 1.000 and camelid 0.000 in both conditions; mouse 0.007 intact and 0.033 framework-masked) but shift toward the middle for AntiBERTy under framework masking (human 1.000 to 0.862; mouse 0.116 to 0.294; camelid 0.000 to 0.541), the behavior expected of a framework-driven call (Fig. 3c). Mask tokens cannot be numbered by ANARCI, so for the five-method comparison we used a numberable substitute: every framework residue was replaced by the cross-species consensus residue at its IMGT position, which removes framework-encoded species information while keeping each sequence a valid antibody chain (Extended Data Fig. 2). IgScope remained the strongest classifier on all four metrics (AUC-ROC 0.930, AUPRC 0.891, accuracy 0.864, MCC 0.697), ahead of AntiBERTy (0.868/0.766/0.742/0.544), whereas the three string- or identity-based scorers lost most of their power (Humatch 0.692/0.475/0.612/0.331; OASis 0.698/0.524/0.597/0.290; T20 0.595/0.469/0.640/0.161). Because the consensus framework is a cross-species mosaic that resembles no natural species, every method loses ground in this condition; consistent with the drop reflecting the foreign framework rather than a loss of CDR signal, IgScope’s decline is concentrated at the human/non-human boundary (at each condition’s balanced-accuracy-optimal threshold, human recall 0.994 to 0.819 and mouse chains called human 0.008 to 0.223), whereas camelid chains are essentially unaffected (0.000 to 0.005).

Consistent with the shortcut interpretation, AntiBERTy’s dominant error is mouse-to-human confusion (44.4% of mouse chains misclassified as human, versus 10.6% for IgScope; Fig. 3e). Finally, on grafting detection, a region-inconsistency score derived from IgScope’s outputs attains an area under the ROC curve of 0.9938 on 69,899 synthetic grafts (Fig. 3d), and in the finer nine-class composition analysis resolves graft direction with MCC 0.9518 versus AntiBERTy’s 6-class 0.7980 (Fig. 3f).

#### Clinical immunogenicity: parity in correlation, superiority in classification

Species labels provide clean ground truth, but the endpoint that ultimately justifies humanness scoring is clinical immunogenicity. We therefore asked whether a multi-resolution score retains the clinical correlation that justifies humanness scoring. On 217 therapeutic antibodies with clinical anti-drug-antibody (ADA) incidence, the IgScope chain score (the arithmetic mean of heavy- and light-chain P(human), which draws on both frameworks and CDRs) correlates with ADA incidence at Pearson r = *−*0.567 (Spearman *ρ* = *−*0.378, R_2_ = 0.321; Fig. 4b), comparable in magnitude to the best published humanness model, HumAb (r = *−* 0.58), and above every other published baseline on this benchmark (Table 1), whether compiled in the BioPhi study ^2^ or reported alongside the recent ImmunoSeq method ^11^ (OASis *−* 0.53; OASis (relaxed) *−* 0.51; germline content *−*0.50; AbLSTM ^23^ *−* 0.47; MG score *−* 0.46; Z-score 0.45; T20*−* 0.41). By Spearman correlation the ordering of the strongest methods shifts, with ImmunoSeq (*ρ* = *−* 0.409) and HumAb ( *−*0.405) marginally above IgScope ( *−*0.378); on this noisy endpoint the two coefficients rank the leading methods differently. The framework-mean score performs equivalently (r = *−*0.569), slightly above a same-protocol recomputation of HumAb (rf-humab package, heavy chain, n = 217: 0.558). Two features derived from the residue track (the longest consecutive stretch with P(human) *<* 0.5, r = +0.568, and the fraction of residues below *−* 0.5, +0.558) correlate at essentially the same level, consistent with non-human *patches*, rather than global dissimilarity, being associated with immunogenicity; framework-region scores ( *−*0.57) are markedly stronger than CDR scores ( *−*0.33), as expected if clinical immunogenicity is driven mainly by framework deviation from human germline.

**Table 1.** Correlation of humanness scores with clinical ADA incidence (n = 217 therapeutics).

| Method | Pearson r | Spearman $\rho$ | R<sup>2</sup> | Source |
| --- | --- | --- | --- | --- |
| AbLSTM<sup>23</sup> | -0.470 | - | 0.221 | Prihoda et al.<sup>2</sup> |
| AbNatiV<sup>4</sup> | -0.488 | -0.356 | 0.238 | Huang et al.<sup>11</sup> |
| AntiBERTy (Fv arithmetic mean) | -0.500 | -0.367 | 0.250 | This work |
| Germline content | -0.500 | - | 0.250 | Prihoda et al.<sup>2</sup> |
| HumAb | -0.580 | - | 0.336 | Prihoda et al.<sup>2</sup> |
| HumAb (same-protocol recomputation) | -0.558 | -0.370 | 0.311 | This work |
| **IgScope chain P(human), arithmetic mean** | **-0.567** | **-0.378** | **0.321** | **This work** |
| ImmunoSeq<sup>11</sup> | -0.530 | -0.409 | 0.281 | Huang et al.<sup>11</sup> |
| MG score | -0.460 | - | 0.212 | Prihoda et al.<sup>2</sup> |
| OASis | -0.530 | - | 0.281 | Prihoda et al.<sup>2</sup> |
| OASis (relaxed)<sup>11</sup> | -0.511 | -0.387 | 0.261 | Huang et al.<sup>11</sup> |
| T20 | -0.410 | - | 0.168 | Prihoda et al.<sup>2</sup> |
| Z-score | -0.450 | - | 0.203 | Prihoda et al.<sup>2</sup> |

**Fig. 4.**
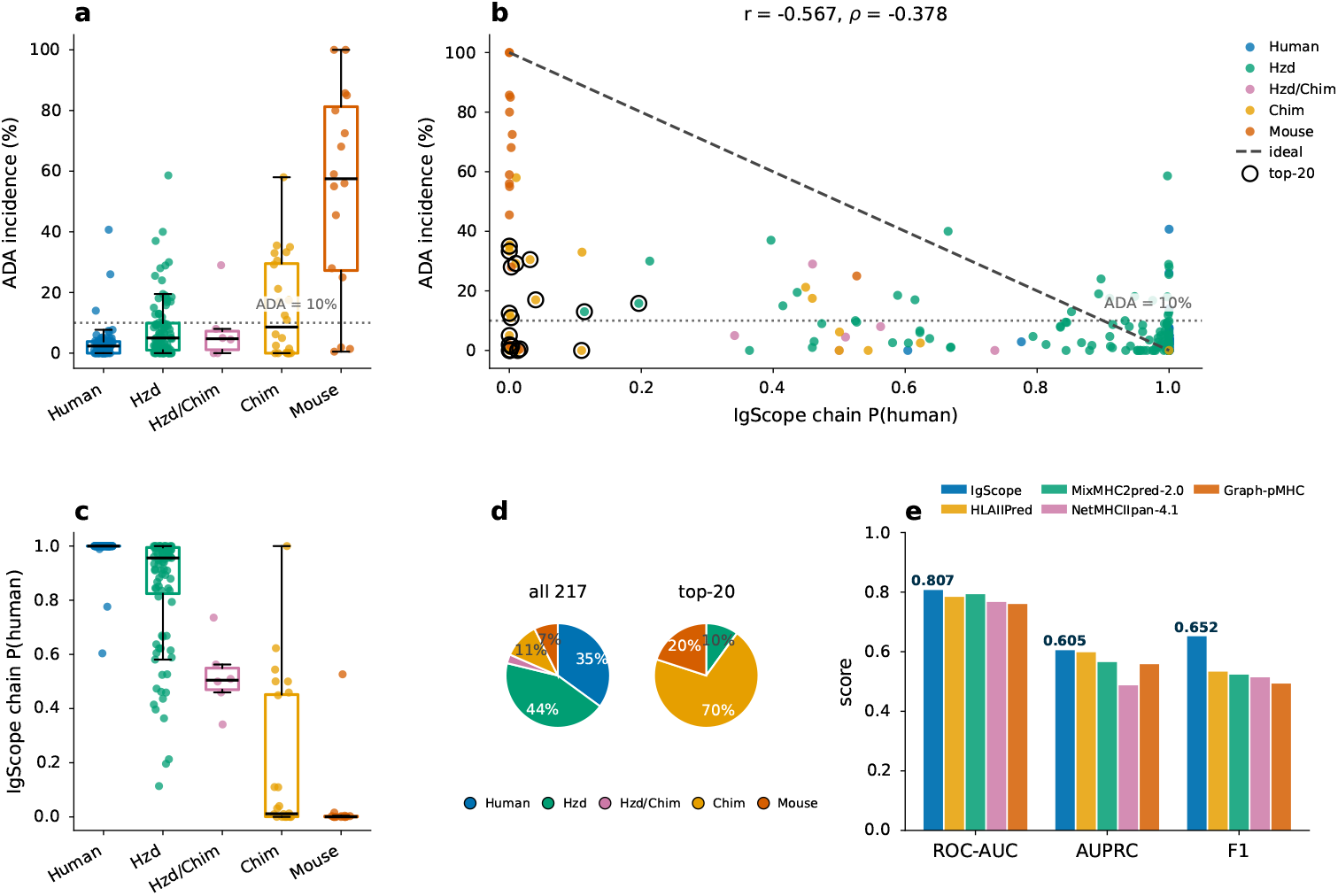
Clinical ADA incidence versus IgScope chain P(human) (n = 217 therapeutics). a, ADA incidence by variable-domain species (dotted line, 10% threshold used in e); boxes, interquartile range; points, individual antibodies. b, ADA incidence against the chain score (arithmetic mean of heavy- and light-chain P(human)), colored by variable-domain species; dashed line, ADA = 100 *×* (1 *−* P(human)); open circles, the twenty largest deviations. r, Pearson correlation; *ρ*, Spearman correlation. c, Chain-score distributions by species. d, Species composition of the full set (left) and of the twenty largest deviations (right). e, Classification of ADA *≤* 10% versus *>* 10% on the 210-antibody deduplicated set, for IgScope and four HLA class II epitope-based predictors; classification threshold at the MCC-optimal chain score (0.46).

Because such scores are used as gates, we then examined where the predictions deviate from the ideal reference line on which ADA equals 100 *×* (1 *−* P(human)), the extreme hypothesis that a fully non-human variable domain elicits ADA in every patient. The twenty antibodies with the largest deviations are overwhelmingly chimeric (14 of 20) or murine (4 of 20), with no human or fully humanized representative, and every deviation lies on the same side of the line: observed ADA far below prediction (Fig. 4b, black circles; Fig. 4d). The per-species score distributions explain the pattern. Variable domains of chimeric or murine origin saturate at P(human) *≈* 0, so the ideal-line prediction collapses onto a single value and cannot discriminate within these classes (Fig. 4c). The saturation is benign for fully murine antibodies, which are indeed highly immunogenic (mean ADA 54%; Fig. 4a); chimeric antibodies (murine variable domains on human constant regions) are clinically well tolerated (mean ADA 14.5%; Fig. 4a), and for them framework humanness has ceased to be the limiting determinant of immunogenicity. As a classifier of ADA *>* 10%, evaluated on the 210-antibody set that remains after removing the five duplicate heavy-light pairs identified in the HLAIIPred study ^12^ (one representative kept per pair on the same side of the threshold; conflicting pairs dropped; 48 of 210 antibodies above threshold, 22.9%), the chain score attains an area under the ROC curve of 0.807 (95% bootstrap confidence interval 0.730–0.876), AUPRC 0.605 and F1 0.652 at the MCC-optimal threshold, exceeding the four HLA class II epitope-based immunogenicity predictors reported on the same cleaned set ^10,12,24–26^ on all three metrics (Fig. 4e; Supplementary Table 3). We read both the correlation and classification results as operating near the ceiling that sequence-only information allows on this noisy endpoint (ADA incidence is confounded by dose, regimen, assay and disease), rather than as a breakthrough.

Literature values are as compiled in the BioPhi study ^2^ or, for AbNatiV, OASis (relaxed) and ImmunoSeq, as reported by Huang et al. ^11^; the HumAb recomputation uses the rf-humab 0.0.10 implementation (heavy chain) on the same 217 antibodies; AntiBERTy values are computed with its native species heads on the same set. All this-work correlations have P *<* 1e-18. Spearman *ρ* was not reported for the BioPhi-compiled baselines. Note that by Spearman correlation the ordering of the strongest methods differs (ImmunoSeq *−*0.409, HumAb *−*0.405, IgScope *−*0.378), consistent with the noise level of this endpoint. The full enumeration of IgScope score definitions is given in Extended Data Table 2.

#### Region-level scores expose non-human CDRs in de novo designed antibodies

We audited 3,359 antibody designs generated de novo by four recent generative models (BoltzGen, IgGM, ODesign and RFantibody; Fabs and VHHs against four targets) against 15,639 natural control chains. In all three chain classes (Fab heavy, Fab light and VHH), every model’s CDRs score significantly below natural CDRs (two-sided Mann–Whitney P *<* 1e-30; Fig. 5a–c), with consistent failure patterns: ODesign is depressed across all CDRs; BoltzGen is bimodal; IgGM fails specifically at CDR1 (median P(human) 0.003) while CDR2/3 are normal, consistent with its template-retention strategy; and RFantibody is lowest across the board, most strikingly for VHHs. Because the ordering of failure modes recurs across chain classes, the defect is a property of the designs rather than of the chain type audited.

**Fig. 5.**
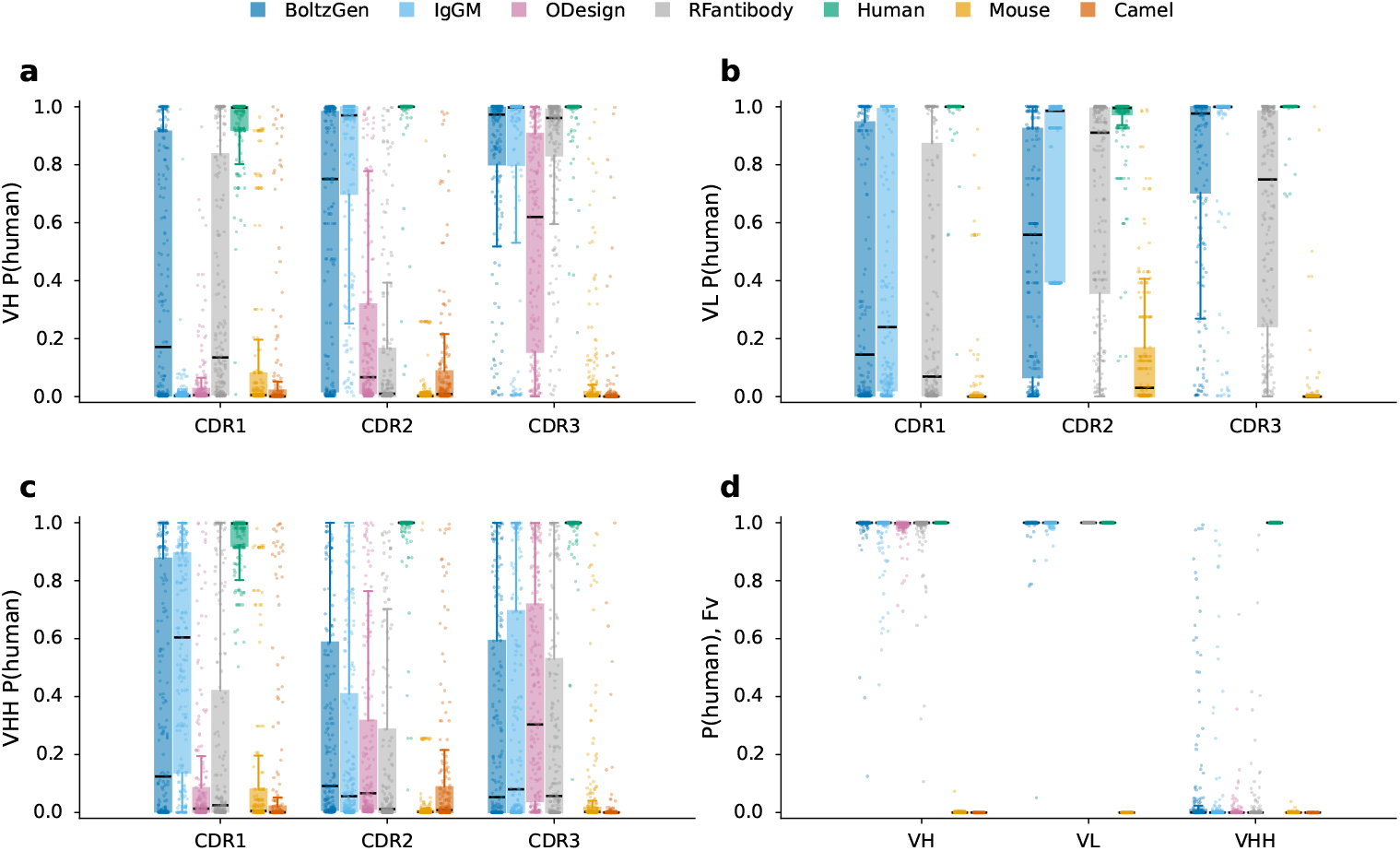
Region-level humanness of de novo designed antibodies. a–c, Per-CDR P(human) 0 for designs from four generative models (BoltzGen, IgGM, ODesign, RFantibody) against natural controls: a, Fab heavy chains (VH); b, Fab light chains (VL); c, VHHs. P(human) is IgScope’s predicted probability that a sequence region is human (0, non-human; 1, human); boxes, interquartile range; points, sampled designs. d, Whole-chain P(human) (Fv) for the same designs by chain class; the design distributions overlap natural human controls.

The decisive observation concerns resolution: the same designs score 0.966–0.988 at the whole-chain level, overlapping natural human controls (Fig. 5d), and the chain-level similarity score OASis under-separates them as well (Spearman *ρ* = 0.561; Supplementary Figure S10). A whole-chain humanness score would pass these designs; among the sequence-level scores tested, only region-level scoring exposes the defect. Because humanness is a routine gate in candidate selection, this blind spot is consequential for pipelines that generate or optimize antibodies computationally.

#### Composition analysis of therapeutics and quality control of humanized-mouse platforms

Applied to 1,095 therapeutic Fabs, IgScope’s composition call (human / humanized / murine / camelid) attains an accuracy of 0.819 against expert annotation (Extended Data Fig. 3). Pairwise reconstruction of the HumAb25 set confirms mechanistic face validity: 31 of 50 humanized chains display the expected framework-human / CDR-murine grafting signature with mouse-to-human V/J family switches, averaging 8.3 framework back-mutation sites per chain (95 of 416 in the vernier zone). As a quality-control application, variable domains from the HIS-Mouse, Kymouse, HUGO-Mab and HUGO-Light platforms score at median *≈*1.0 across the full domain and all seven regions (Fig. 6a, b; not every platform contributes every chain class, and there are no Kymouse or HUGO-Light light chains). HUGO-Nano VHHs also score as human at the chain level (median Fv P(human) 1.000; no chain below 0.9), with one biologically expected exception at the region level: their FR2 is bimodal (Fig. 6c). The 25.2% of chains that carry the camelid hallmark tetrad FERF at IMGT 42, 49, 50 and 52 have a median FR2 P(human) of 0.001 (100% of chains *<* 0.5), whereas the remaining 74.8% score 0.999 (97.9% *>* 0.9). FERF is engineered onto human V/D/J genes and required for VHH stability. Masking only these four positions in the FERF chains restores the FR2 score (median 0.989; 99.8% *>* 0.9), whereas masking four random FR2 positions leaves it at 0.0003; the other six region scores are unchanged (Δ = 0 to numerical precision) and the chain score stays at 1 in all three conditions (Fig. 6d). The departure is equally confined at the sequence level: at the four hallmark positions HUGO-Nano’s most frequent residue is the human one, with a 26–27% camelid minority (V42/F, G49/E, L50/R, W52/F) whose camelid member occurs in at most 0.5% of human VH chains, whereas at 11 of the other 13 FR2 positions its most frequent residue is also the most frequent human one, the remaining two (IMGT 54, 55) carrying residues present in 31% and 10% of human VH chains (Fig. 6e). The non-human signal is therefore a four-residue design choice confined to the camelized subpopulation, not a failure of humanization, and it is precisely the localized, intended non-humanness that a chain-level score cannot resolve. It is visible under both calibrations of the region score (Supplementary Fig. S6).

**Fig. 6.**
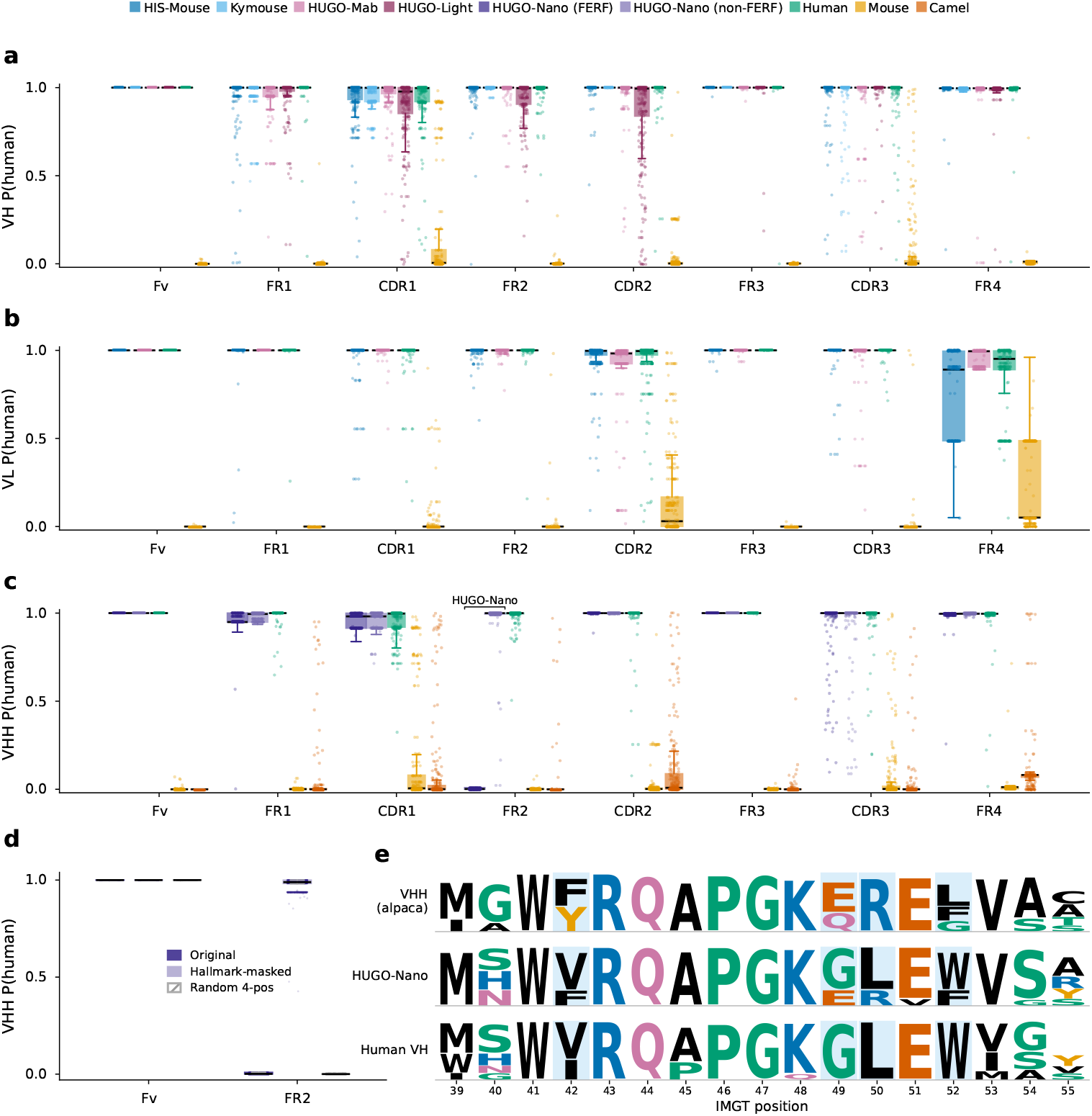
Humanized-mouse platforms produce humanized variable domains, with one intended exception. a, VH from HIS-Mouse (10,000), Kymouse (10,000), HUGO-Mab (915) and HUGO-Light (1,472), against natural human and mouse VH; b, VL from HIS-Mouse (20,000) and HUGO-Mab (915), same references; c, HUGO-Nano VHHs (10,000), against natural human, mouse and camel VHHs; one platform, decomposed by the engineered camelid FERF tetrad (IMGT 42, 49, 50, 52; 25.2%), the remaining non-FERF chains carrying human-type tetrads. P(human) (Fig. 5) for the variable domain and seven IMGT regions; boxes, interquartile range; points, sampled sequences. d, HUGO-Nano FERF chains unmodified, hallmark-masked or random-masked: masking the four raises median FR2 from 0.001 to 0.989, the random mask leaving it at 0.0003; chain scores remain *≈* 1. e, FR2 composition (IMGT 39–55) of alpaca VHHs, HUGO-Nano and human VH; letter height, residue frequency (*<*10% omitted).

#### The excess of sub-0.9 CDR scores in HUGO-Light is compositional

Fig. 6a shows that HUGO-Light’s VH CDR1 and CDR2 carry a much larger share of scores below 0.9 than any other platform. Because a region score depends only on the scored region’s own residues (Methods), that excess must come from the sequences themselves, and germline composition accounts for it. In both regions HUGO-Light’s CDRs are the furthest from their own human germline allele (identity 27.1% in CDR1 and 23.3% in CDR2, versus 41.3–100% elsewhere) and carry 1.6–2.7-fold more substitutions per residue (0.158 and 0.190); they therefore carry the largest share of chains in the ambiguous 0.5–0.9 band (23.3% and 21.5%, versus at most 12.3% and 6.2%). The relation is graded within platforms as well as across them—mutated peptides score below germline-identical ones wherever mutations exist (Cliff’s *δ* = 0.61–0.69 in CDR2)—so sub-0.9 scores are the expected output for germline-distant CDRs. Exact sharing of CDR peptides between human and mouse germlines, the obvious alternative source of intermediate scores, is rarest in HUGO-Light (8.9% of CDR1 chains versus 17.8–33.4%) and can account for at most 4.6 of its 23.3 percentage points. Because the cohorts also differ in provenance and immunization status (the Kymouse repertoire is entirely unmutated), we read the comparison as sequence composition, not as a platform ranking.

#### Proof-of-concept clonal lineage reconstruction

Finally, as an application of the evolution-aware heads, IgScope’s predicted germlines and a language-model edge weight (parent negative log-likelihood) orient clonal lineage trees by minimum directed spanning trees (Methods). Across 6,966 clonotypes derived from 20.9 million repertoire sequences, somatic hypermutation accumulates monotonically along root-to-tip paths (monotonicity 0.979, versus 0.951 for Hamming-distance trees; Extended Data Fig. 4), and in a 30-member affinity-maturation series tree depth does not correlate significantly with binding affinity under the final checkpoint (r = *−*0.319, P = 0.086; Extended Data Fig. 5). We therefore present lineage reconstruction as a proof of concept, not a validated affinity predictor.

## Discussion

The through-line of this study is that the *resolution* of a humanness score determines what it can see. Whole-chain scores saturate: IgScope reaches 0.9998 internal accuracy and ADA correlations near r *≈ −*0.57, like the best published whole-chain methods. Yet they pass de novo designs whose CDRs are measurably non-human in all three chain classes audited, and miss four camelid hallmark residues in the FR2 of otherwise fully humanized chimeric VHHs. Fine-grained scores are meaningful only if they are local, and locality does not come for free: our framework-masking and chimera experiments show that it must be engineered through architecture and supervision, and measured rather than assumed. We suggest that the framework-masked CDR test become a standard diagnostic for any model claiming region-level interpretability.

Region-resolved scoring sits naturally alongside existing humanness methods, which output chain-level (occasionally V-gene-level) scores and do not audit whether CDR signal is used ^2,4,5,27,28^. Among antibody language models ^13,29–34^, only AntiBERTy provides native classification heads, and none provides region-isolated representations or region-level species supervision; general protein language models, though widely applied to antibodies, offer no antibody-specific supervision at any resolution ^35,36^; evolution-aware pretraining objectives are shared in spirit with EATLM ^31^, but in IgScope they are tied to a verified-locality design rather than used as a general representation-learning signal.

Several limitations bound the interpretation of our results. First, region-level accuracy is limited by shared germline motifs: when a CDR sequence is identical between human and mouse germline families, no region-local model can beat chance on that instance, and the reported accuracies are close to this sequence-determined ceiling. Second, the lineage analysis is checkpoint-sensitive and lacks simulated-repertoire benchmarks against established phylogenetics tools ^37–39^. Third, the scorer head-to-head (Fig. 3a, b) excludes AbNatiV, which provides separate nativeness models per chain type rather than a single cross-species humanness score, and uses a 30,000-chain stratified subsample because several scorers do not scale to the full external set. Fourth, the model scores single chains independently; paired-chain effects, VH–VL interfaces and structure are outside its current scope. Finally, the ADA endpoint itself is noisy, and r *≈ −*0.57 may be near the sequence-only ceiling of that benchmark.

Beyond antibody engineering, the pattern we expose (chain-level metrics that conceal localized defects) is likely to recur wherever generative models are screened by similarity scores. Coupling generators to region-level audits is a direct next step, as is coupling the residue track to structure-aware humanization, for example with generative humanization tools ^40^.

## Methods

### Data

Training used the full dev data corpus of 119M chains, assembled from two sources: the Observed Antibody Space ^1^ and, because OAS contains comparatively few camelid sequences, an alpaca VHH repertoire dataset ^41^ (pre-immunization PBMCs from 22 alpacas, sequenced with VHH libraries built from hinge- and leader-region primers; 7.5M chains after cleaning). Both sources were filtered to remove non-functional, frameshift and premature-stop records, to CDR-defined lengths of 80–148 amino acids, enforcing conserved cysteines, allowing at most two ambiguous (X) residues, and de-duplicating within species. Redundancy was further reduced by CDR3-grouped Linclust clustering at 95% identity ^42^. The split was stratified by species *×* chain type at 95/2/3 (train/validation/test); the test set contains 2,379,517 chains (Supplementary Table 1). The camelid class pools camel and alpaca sequences, so species-level conclusions for camelids should be read at the family level. Sequences from humanized-mouse platforms (Kymouse ^43^; HIS-Mouse) were excluded from training entirely.

### Mutation-state labels

The Observed Antibody Space provides, for each sequence, a per-position germline alignment. From this alignment we derive two binary position labels. A somatic-hypermutation (SHM) label is set to 1 at positions where the germline residue is a standard amino acid and differs from the observed residue; a non-templated-insertion (NP) label is set to 1 at positions whose germline column is non-amino-acid (X or gap), corresponding to V(D)J junctional insertions; positions that are neither form the germline-encoded class. These labels have three uses: they define the three-way mutation-process prediction (MPP) target at every position; they mark the positions at which the ancestral-germline-prediction (AGP) target is replaced by a dedicated [GAP] token; and their per-sequence frequencies are the SHM and NP rates reported in Supplementary Table 1 and used to group positions in all mutation-state-resolved analyses.

### Chimeric and back-mutation augmentation

Two chimera types were constructed from same-locus donors of different species: a V-block swap exchanging IMGT positions 1–104 (FR1–CDR3) against CDR3+FR4, and a free region-set swap, each with 50% probability. Region and residue labels on chimeras equal the donor species of each segment, and the chain-level loss is ignored on chimeras. Chimeras constitute 30% of the training stream. A second augmentation replaces 1–5 framework positions with single donor-species residues, with the replaced positions carrying the donor-species residue label. Augmentations were applied to the training partition only; validation and test sets remain purely natural.

### External evaluation sets

All external sets are disjoint from the training partition. Species classification and grafting detection use the VDJServer external test set (Supplementary Table 2), 1,005,481 BCR chains curated from six studies deposited in the VDJServer AIRR Data Commons ^15–22^; none of these studies contributes to the training corpus. VDJServer is a public AIRR Data Commons that contained 39 studies, 3,408 repertoires and approximately 2.5 billion annotated rearrangements at its December 2021 release; we retained its BCR (IGH, IGK and IGL) rearrangements and discarded the TCR records. Clinical immunogenicity is benchmarked on ADA217, 217 therapeutic antibodies with clinically measured ADA incidence curated by Hu-mAb and BioPhi ^2,5^; the binary classification task uses the 210-antibody subset obtained after removing duplicate heavy–light pairs, following Haghighatlari et al. ^12^. Composition analysis uses 1,095 therapeutic Fabs of Thera-SAbDab ^44^ with expert annotations (human, humanized, murine, camelid). The de novo design audit scores 3,359 designs from four generative models (BoltzGen, IgGM, ODesign and RFantibody ^6–9^; Fabs and VHHs against four targets) alongside 15,639 natural control chains. Quality control uses repertoires of the humanized-mouse platforms HIS-Mouse, Kymouse, HUGO-Mab, HUGO-Light and HUGO-Nano. Clonal lineage reconstruction uses a repertoire sequencing dataset of 20.9 million sequences, from which 6,966 clonotypes are built. Transfer benchmarks comprise the SAbDab paratope split ^45^ and the ATUE benchmark ^31^.

### Germline and score analysis of humanized-mouse platforms

The cross-platform comparison (Extended Data Table 3) uses VH chains only, de-duplicated on the aligned sequence and restricted to chains carrying region-head scores, i.e. the cohort of Fig. 6a; HUGO-Mab contributes 581 unique VH sequences from 915 antibodies. CDR1 (IMGT 27–38) and CDR2 (56–65) peptides were cut from the aligned sequence using the ANARCI numbering. Each chain was matched to a human germline allele through its v call field; IMGT duplicated alleles (the ‘d’ suffix, e.g. IGHV3-20*d01) were mapped to the corresponding numbered allele, and the 0.03–0.3% of chains whose allele is absent from the reference set were excluded from the germline-identity columns. A CDR was called germline-identical when its peptide equaled that allele’s CDR, and the substitution rate is the number of mismatched positions per CDR residue. A peptide was called shared when it occurred in at least one human and one mouse germline allele (Supplementary Data 1). The ambiguous band is 0.5 *≤ P* (human) *≤* 0.9; Cliff’s *δ* is computed from the Mann–Whitney *U* statistic for germline-identical versus mutated peptides and is left undefined where a platform has no mutated chains.

### Counterfactual masking of the HUGO-Nano FR2

The 10,000 HUGO-Nano VHH chains of Fig. 6c (sampled from 99,690 processed chains) were scored three times: unmodified; with the residues at IMGT positions 42, 49, 50 and 52 replaced by the mask token; and with four non-hallmark FR2 positions (IMGT 39–55) drawn at random per chain and masked. Because Module A attends only within a region and the region head pools only within-region states, only the FR2 score can change, and the other six region scores differed from the unmodified run by exactly 0. Fig. 6d is restricted to the 2,517 chains carrying FERF at all four positions; the remaining 7,483 chains carry human-type tetrads and score *≈* 1 in all three conditions. The FR2 logo (Fig. 6e) covers the full FR2-IMGT, positions 39–55, computed over up to 10,000 ANARCI-numbered chains per source (natural alpaca VHHs and human VH from the held-out partitions, HUGO-Nano from the platform repertoire); letters are drawn for residues present in at least 10% of chains at that position.

### Model architecture

IgScope is a single-chain (VH/VL/VHH) Transformer encoder. Input chains are IMGT-numbered ^46,47^, truncated to a maximum of 168 tokens including class and end tokens, and embedded as the sum of three tables: a residue embedding (vocabulary of 33, including a ‘[GAP]’ token and mask/special tokens), a factorized IMGT position embedding (learned tables over the 130 numeric IMGT positions and over 9 insertion codes) and a region embedding over the seven IMGT regions. Module A comprises four Transformer layers (d model 480, eight heads, feed-forward dimension 1,920, dropout 0.1) with a same-region attention mask; all seven regions share one weight set. A masked mean-pool over each region yields a [7 *×* D] readout, scored by a RegionHead shared across regions. Because attention is confined to same-region pairs and the readout averages only within-region states, a region score is a function of that region’s own residues: an identical CDR peptide receives an identical score in every repertoire, so cross-repertoire differences in region scores reflect sequence composition rather than model state. Module B comprises four standard Transformer layers over the full sequence; its class token feeds the ChainHead (three-way species) and auxiliary chain-type, V-gene and J-gene heads, and its per-token states feed the MLMHead and a shared two-layer MLP+GELU+dropout evolution trunk, from which the ResidueHead (per-residue species), AGPHead (germline reconstruction, with ‘[GAP]’ marking non-templated insertions) and MPPHead (three-way germline / somatic-hypermutation / non-templated) are computed. Species, chain type and V/J genes are never inputs. Total parameters: 23.0M.

### Training

The training objective is a weighted sum of nine cross-entropy terms,

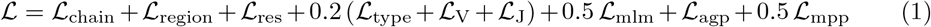

in which the chain, region and residue terms carry unit weight, the auxiliary chain-type, V-gene and J-gene terms 0.2 each, the masked-language-modeling term 0.5, and the AGP and MPP terms 1.0 and 0.5, respectively. The chain loss ignores chimeras, whose chain-level label is intentionally undefined. The region loss is the weighted mean of the seven per-region cross-entropies, with CDR regions weighted twice as heavily as framework regions. The residue, AGP and MPP losses use focal loss ^48^,

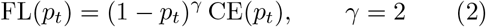

which de-emphasizes the easily predicted germline-encoded majority at which an unweighted loss saturates. Masking followed Ab-ModMask ^32^: each batch draws uniformly at random one of three modes — independent per-position masking, short segments of 3–5 tokens, or a single long segment — with the masking rate *ρ* sampled from [0.10, 0.40]; the segment modes mask approximately *ρ*_·_n residues, where n is the number of residues in the sequence. In all modes, candidate positions are sampled in proportion to their weight w, the per-position entropy of the training-set amino-acid distribution,

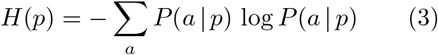

normalized within each sequence to mean one, so that hypervariable positions (the CDRs) are masked more frequently. Optimization used AdamW ^49^ with learning rate 1e-4, weight decay 0.01, 5% warmup, a cosine schedule, gradient clipping 1.0 and automatic mixed precision; the global batch was 768 (384 2 *×* GPUs, distributed data parallel, PyTorch 2.4.1), with 10 epochs planned, seed 42 and early-stopping patience 3. The best checkpoint (epoch 1, step 290,001) was selected by validation CDR-mean area under the precision–recall curve (0.9975).

Masked-language-modeling performance is reported as held-out perplexity over a residue subset *M* —all residues, germline-encoded residues, somatic-hypermutation sites or non-templated insertions—with every residue in *M* replaced by a mask token and all other positions left visible:

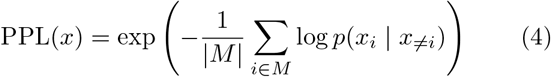

where |*M*| is the number of scored positions, *p*(*x*_*i*_ | *x*_*≠i*_) is the model probability of the true residue at position *i* given all unmasked positions, and the test-set value is the arithmetic mean of PPL(*x*) over chains, so that per-sequence perplexities are averaged after, not before, exponentiation ^32^.

### Evaluation suites

Internal diagnostics were computed on the full held-out test set with the best checkpoint and include the framework-masked CDR test (all framework tokens replaced with mask tokens before re-scoring CDRs), the back-mutation diagnostic (synthetic murine-to-human framework reintroductions), chimera region accuracy and chain-level P(human) propagation, shared-motif attribution (IMGT-annotated human–mouse shared CDR motifs), and per-residue and auxiliary-task metrics (Extended Data Table 1). Error enrichment on shared motifs is reported as an odds ratio,

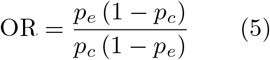

where *p*_*e*_ is the fraction of species-s CDR errors that fall on shared motifs and *p*_*c*_ the corresponding fraction among correct calls. For graft-versus-natural detection (Fig. 3d), each chain receives a region-inconsistency score

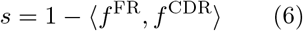

where *f* ^FR^ and *f* ^CDR^ are three-component vectors indexed by species (human, mouse, camelid): component *k* is the mean predicted probability of species *k* over the four framework regions (*f* ^FR^) or the three CDR regions (*f* ^CDR^); *⟨·, ·⟩* is the standard inner product of these two vectors (the angle brackets are not a sequence interval), so *s* is one minus their overlap and approaches 1 when the framework- and CDR-averaged species distributions disagree. The ROC analysis compared 69,899 synthetic grafts with a random sample of 50,000 natural chains. ADA correlations report Pearson r, Spearman *ρ*, R_2_ and P values on n = 217 therapeutics, with the chain score defined as the arithmetic mean of heavy- and light-chain P(human) (all aggregation calibers are enumerated in Extended Data Table 2); values for ImmunoSeq, AbNatiV and OASis (relaxed) are as reported by Huang et al. ^11^; the HumAb baseline was recomputed with the rf-humab package under the same protocol. Binary classification of ADA *≤* 10% versus *>* 10% followed the deduplicated 210-antibody protocol of Haghighatlari et al. ^12^ (five duplicate heavy-light pairs removed: one representative kept per pair on the same side of the threshold, conflicting pairs removed entirely); the IgScope F1 score is reported at the MCC-optimal chain-score threshold (0.46), with 2,000-replicate bootstrap confidence intervals for ROC-AUC and AUPRC. VDJServer evaluation used bootstrap standard deviations (200 replicates). For the humanness-scorer comparison (Fig. 3a, b), a stratified subsample of the external natural set (10,000 chains per species, seed 42) was scored with OASis (the promb implementation of the OASis 9-mer, greater-than-10%-of-subjects peptide set), T20 (mean BLASTp identity to the 20 closest human VH sequences; Llamanade implementation), Humatch (heavy and light classifiers; humanness taken as one minus the non-human class probability) and the AntiBERTy native species head; IgScope scores were recomputed on the same chains. For display in Fig. 3a, each method’s native score was min–max rescaled to [0, 1] across all chains shown, per method rather than per species: IgScope, Humatch and AntiBERTy output probabilities, OASis reports identity to the nearest human reference sequence (observed range 0.065–1.000), and T20 reports a 0–100 score (observed range 42.3–99.9; VH chains). Species-discrimination metrics (AUC-ROC and AUPRC) were computed with scikit-learn, with human as the positive class. The framework ablations used a 30,000-chain benchmark drawn from the same external VDJServer repertoires (10,000 chains per species; 29,985 heavy and 15 kappa chains), with the identical chains scored under every condition. For the consensus-framework ablation (Extended Data Fig. 2), every framework residue of every chain in the 30,000-chain benchmark was replaced by the modal residue at the same IMGT position (number and insertion code) across all benchmark chains of the same chain type, leaving CDRs untouched; this makes the framework identical across species while keeping the sequence a valid, numberable antibody chain. Accuracy and MCC use each method’s balanced-accuracy-optimal (Youden) threshold, recorded in the source data; the mask-token ablations restricted to the two language models (either every token in FR1–FR4 or every token in CDR1–CDR3 replaced by that model’s mask token—<mask> for IgScope and [MASK] for AntiBERTy—with the complementary regions untouched and the same chains scored) are shown in Fig. 3c.

Lineage reconstruction used AGP-predicted germlines as roots, parent negative log-likelihood edge weights and minimum directed spanning trees, evaluated by SHM monotonicity along root-to-tip paths.

## Data availability

The training corpus was assembled from the Observed Antibody Space ^1^ and the alpaca VHH repertoire dataset of Wang et al. ^41^; the OAS component is available from the OAS authors under their data-use terms and the alpaca component from the original publication. The cleaning, clustering and split pipelines are included in our code release. VDJServer data are available from the VDJServer AIRR Data Commons ^21,22^ under their access policy. ADA incidence for the 217 therapeutics follows the Hu-mAb/BioPhi benchmark ^2,5^. Thera-SAbDab is available at its public database ^44^. Per-V-gene human–mouse germline CDR similarity tables accompany this paper as Supplementary Data 1. De novo design sets were generated with the respective models (BoltzGen, IgGM, ODesign, RFantibody). Source data for all figures will be deposited with the manuscript. All other data are available from the corresponding author on reasonable request.

## Code availability

The IgScope source code, model configurations and evaluation scripts have been deposited in a private repository at [https://github.com/Huatsing-Lau/IgScope](https://github.com/Huatsing-Lau/IgScope); they will be made publicly available, with persistent archiving (Zenodo DOI), upon publication of this article. In the meantime, the code is available to editors and reviewers on request to the corresponding authors. Checkpoints for the main results and for the lineage case study will be deposited together with the code release.

## Acknowledgements

We thank the Macao Science and Technology Development Fund (No. FDCT 0019/2024/RIA1) for funding this research. We thank Yuxiang Liu and Xing Mou (Cyagen Biomodels (Guangzhou)) for providing the NGS data of HUGO-Nano mice, and Lunji Wang (Artificial Intelligence Innovation Center, Research Institute of Tsinghua, Pearl River Delta) for GPU server support.

## Author contributions

H.L.: Conceptualization, methodology, software, investigation, data curation, formal analysis, validation, visualization, writing—original draft, writing—review & editing, resources. P.C.: Conceptualization, methodology, software, investigation. Shuxian Zhou: Data curation, investigation. B.W.: Data curation, validation. S.D.: Validation. L.Z.: Validation. X.L.: Writing—review & editing. Shun Zhou: Conceptualization, resources. L.H.: Conceptualization, supervision, project administration, resources, funding acquisition. L.Y.: Conceptualization, formal analysis, resources, supervision, funding acquisition, writing—review & editing.

## Competing interests

L.H. is a founder of Cyagen Biomodels (Guangzhou). The HUGO-Mab, HUGO-Light and HUGO-Nano mice evaluated in this study are proprietary products of Cyagen Biomodels. The other authors declare no competing interests.

## Additional information

**Supplementary information** is available for this paper.

## Extended Data

**Extended Data Fig. 1.**
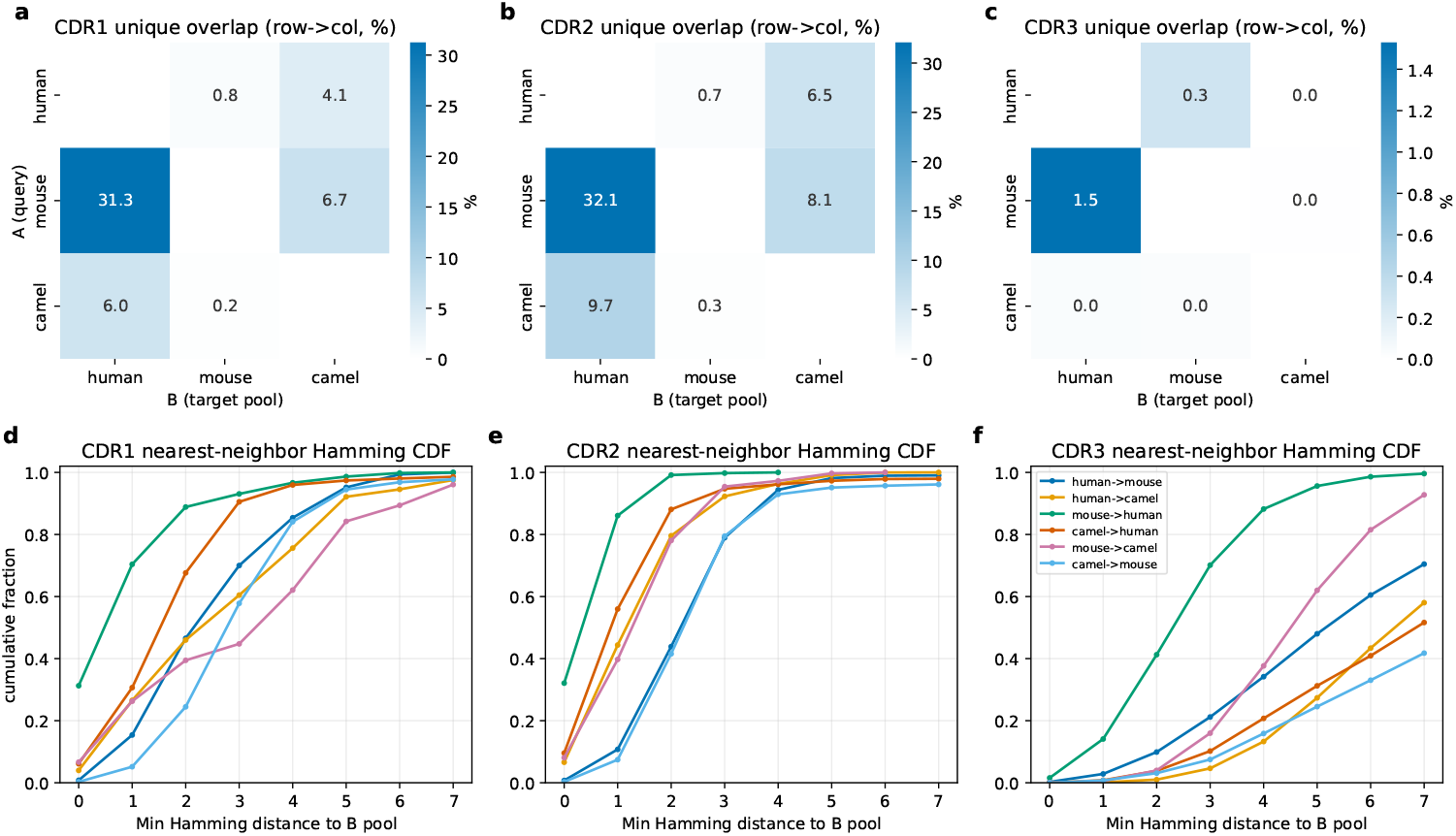
Cross-species CDR overlap explains the region-accuracy ordering of Fig. 2. a-c, Unique cross-species CDR overlap rates (percentage of species-A unique CDRs found verbatim in species-B pool; row *→* column) for CDR1 (a), CDR2 (b) and CDR3 (c). d-f, Cumulative distribution of the nearest-neighbor Hamming distance from each species-A unique CDR to the 5 same-length pool of species-B unique CDRs. Murine CDR1/CDR2 are frequently identical or near-identical to human CDRs (mouse *→* human unique overlap 31.3% / 32.1%; 70.4% / 86.1% within one substitution), whereas CDR3 shows essentially no cross-species overlap (unique ¡=1.5%, median Hamming distance 3-7). This information asymmetry is the sequence-level reason why region-level accuracy on CDR1 (0.9618) and CDR2 (0.9731) lags CDR3 (0.9771) and the framework regions, and 9 underlies the shared-motif attribution of Fig. 2e, f. Analysis on the held-out validation and test pools (4,759,029 chains) after Linclust 95% de-duplication. Per-V-gene CDR1/CDR2 sequences and pair-wise identities are provided in Supplementary Data 1.

**Extended Data Fig. 2.**
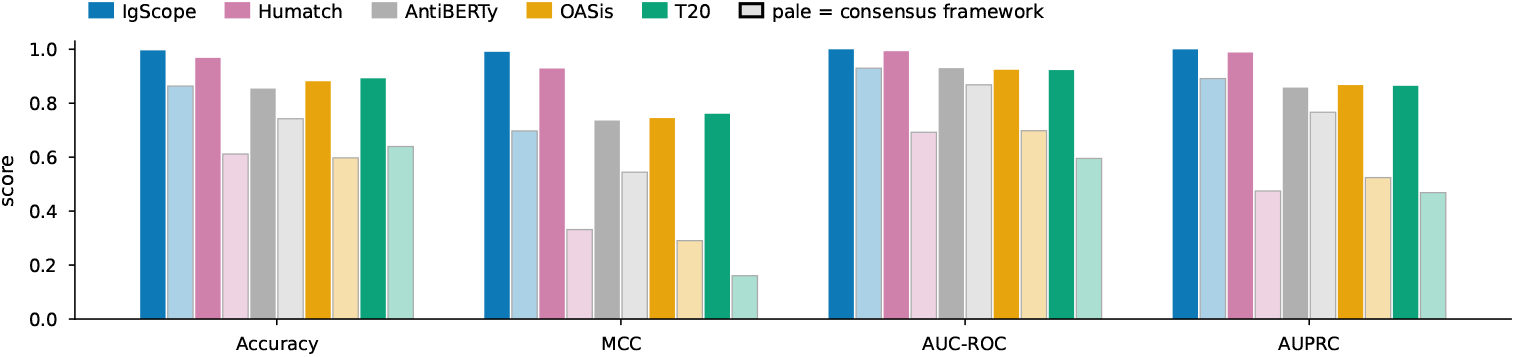
Consensus-framework ablation across all five scorers. Binary species classification (human versus non-human) on the 30,000-chain benchmark (10,000 chains per species) with intact sequences (solid bars) and with every framework residue replaced by the cross-species consensus residue at its IMGT position (pale bars); accuracy and MCC are evaluated at each method’s balanced-accuracy-optimal threshold. The consensus framework leaves every input a numberable antibody chain, which is why the string- and identity-based scorers can be included, but because it is a cross-species mosaic resembling no natural species, every method loses ground, so the condition is a stress test rather than a clean measure of framework reliance. IgScope remains strongest on all four metrics (AUC-ROC 0.930, AUPRC 0.891, accuracy 0.864, MCC 0.697), ahead of AntiBERTy (0.868/0.766/0.742/0.544), whereas Humatch (0.692/0.475/0.612/0.331), OASis (0.698/0.524/0.597/0.290) and T20 (0.595/0.469/0.640/0.161) lose most of their power. The framework mask-token ablation of Fig. 3c, which removes framework content without introducing a foreign scaffold, is the causal test for the two language models.

**Extended Data Fig. 3.**
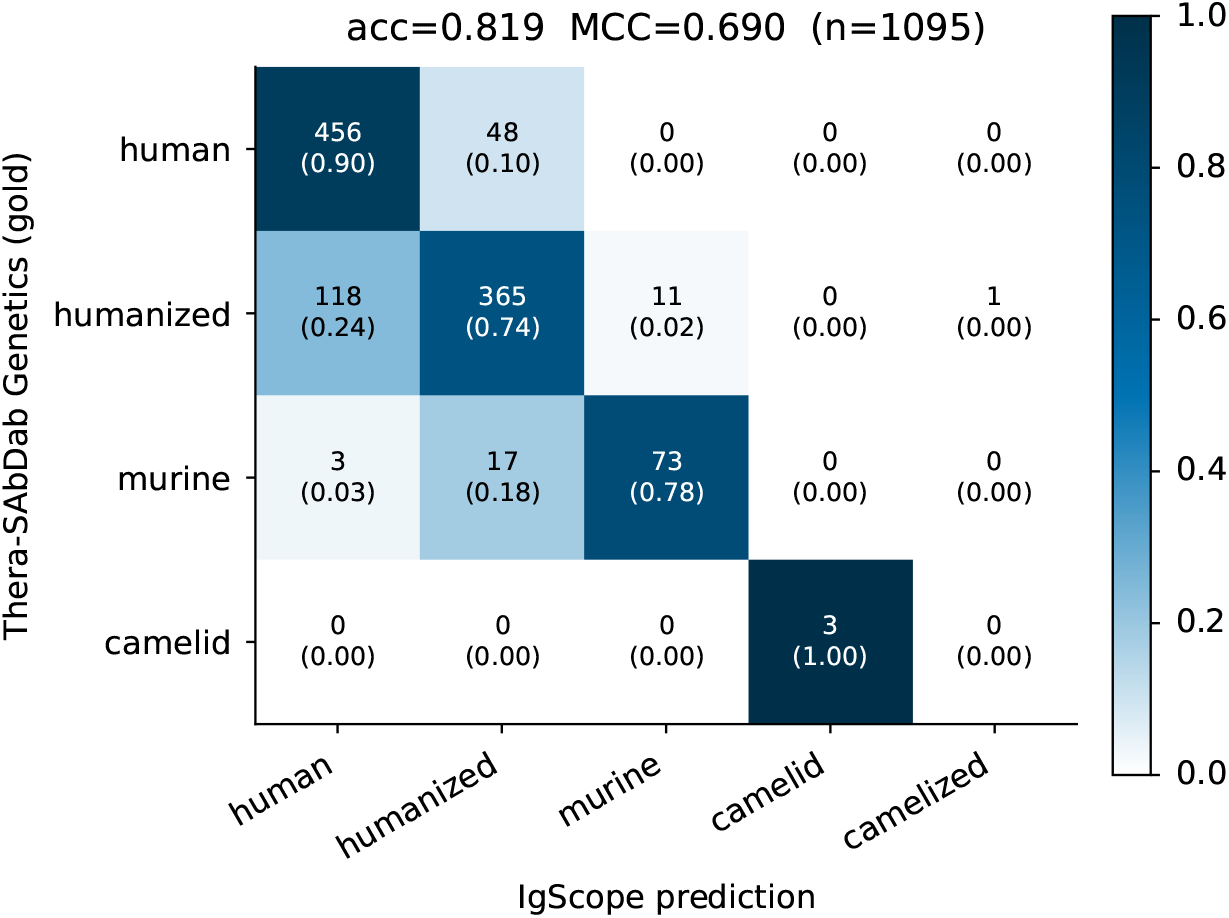
Thera-SAbDab composition classification (n = 1,095 therapeutic Fabs). Confusion matrix of IgScope composition calls (human / humanized / murine / camelid) against expert annotation (accuracy 0.819, MCC 0.690); the dominant confusion is humanized *→* human (118 cases) — sequences for which the distinction is essentially impossible.

**Extended Data Fig. 4.**
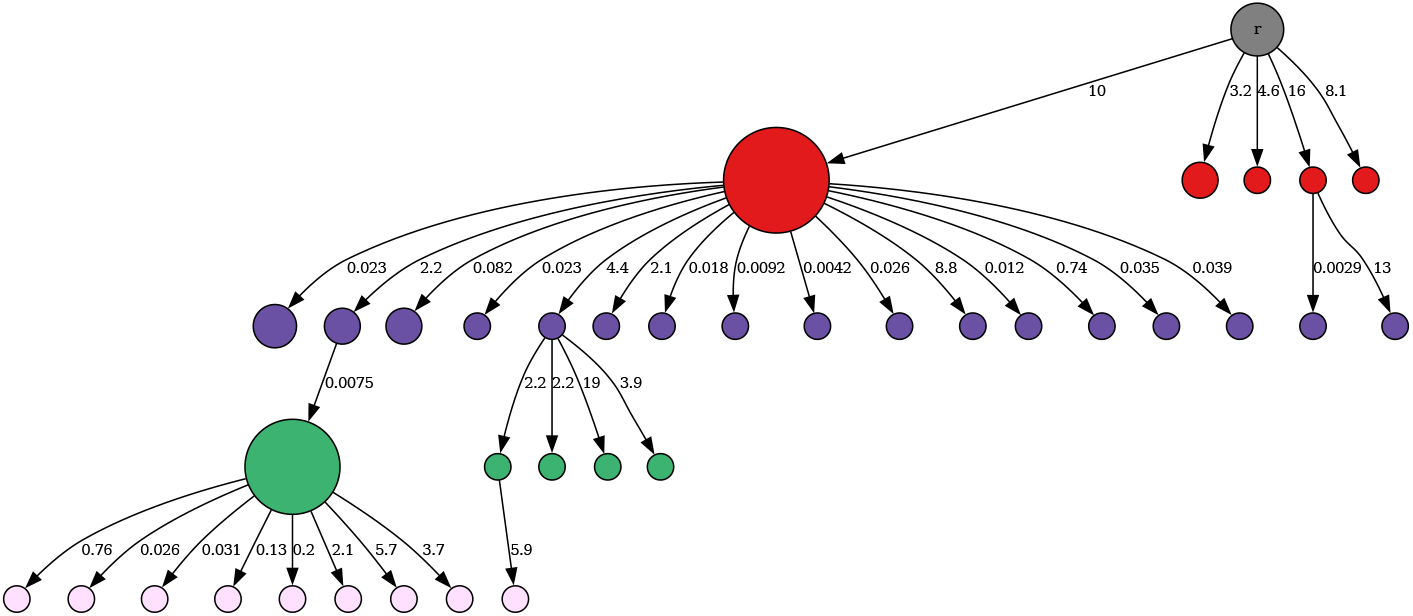
Example reconstructed lineage (T015 repertoire, clonotype 1 IGHV4-59-02/IGHJ5-02). AGP-predicted germline root; edges weighted by parent-NLL; tree by 2 minimum directed arborescence. SHM accumulates monotonically along root-to-tip paths (monotonicity 0.979 across the repertoire).

**Extended Data Fig. 5.**
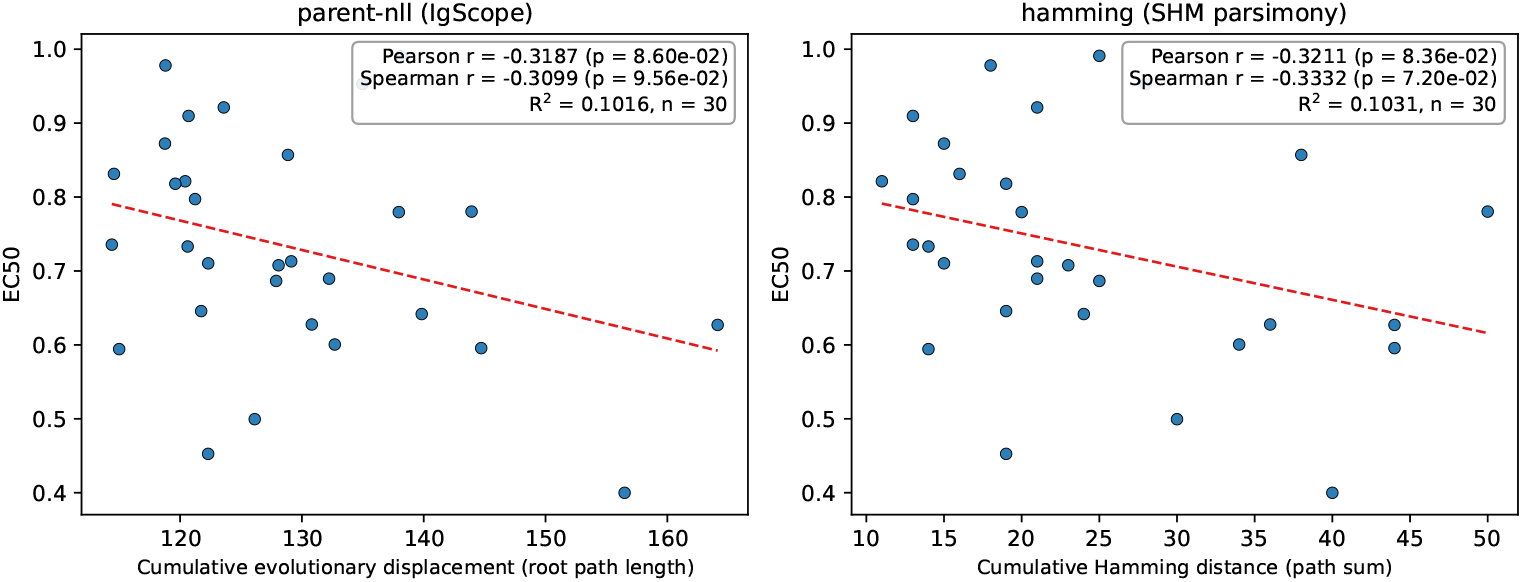
Depth–EC50 correlation under two tree constructions (final 2 model checkpoint, n = 30). Left: parent-NLL tree — cumulative evolutionary displacement 3 (root path length) versus EC50, Pearson r = *−*0.319 (P = 0.086). Right: Hamming-distance (SHM-parsimony) tree — cumulative Hamming distance (path sum) versus EC50, r = *−* 0.321 (P = 0.084); this baseline uses no model and is therefore checkpoint-independent. Neither reaches significance at n = 30; the negative trend is consistent across tree constructions.

**Extended Data Table 1.** Internal diagnostic suite master table (test set n = 2,379,517; V- and J-family accuracies from the validation set).

| Diagnostic | Metric | Value |
| --- | --- | --- |
| Chain type (H/K/L) | acc | 1.0000 |
| V family | acc | 0.9998 |
| J family | acc | 0.9853 |
| Chain-level species | acc / macro-F1 | 0.9998 / 0.9997 |
|  | AUC-ROC / AUPRC | 0.99999 / 1.00000 |
| Region-level species | CDR1/2/3 acc | 0.9618 / 0.9731 / 0.9771 |
| by chain type | CDR/FR mean, H (2.08M) | 0.9695 / 0.9944 |
|  | CDR/FR mean, K (0.16M) | 0.9675 / 0.9972 |
|  | CDR/FR mean, L (0.14M) | 0.9916 / 0.8998 |
| Chimera (3 pairs, n=6,000) | CDR mean / FR mean acc | 0.941 / 0.980 |
| Unique CDR species | exclusive acc CDR1/2/3 | 0.9528 / 0.9022 / 0.9777 |
|  | shared consistent-rate CDR1/2/3 | 0.9988 / 0.9969 / 0.9981 |
| Residue-level species | AUPRC all/SHM/NP/germline | 0.999997/0.999987/0.999998/0.999998 |
|  | ROC-AUC (same order) | 0.999989/0.999969/0.999993/0.999991 |
| Back-mutation (vernier) | backmut acc | 0.9338 |
|  | vernier unmut. / FR other / CDR | 0.9974 / 0.9923 / 0.9876 |
| AGP | token acc | 0.9925 |
|  | [GAP] recall / F1 / AUPRC | 0.926 / 0.925 / 0.981 |
| MPP | 3-class acc / macro-F1 | 0.9927 / 0.966 |
|  | SHM recall / F1; NP recall / F1 | 0.969 / 0.975; 0.923 / 0.925 |
| MLM | acc | 0.868 |
| MLM | ppl all / germline / SHM / NP | 2.63 / 2.06 / 1429 / 20.8 |

**Extended Data Table 2.** Complete enumeration of score definitions and baselines for the ADA correlation (n = 217 therapeutics).

| Method | Pearson r | Spearman $\rho$ | R<sup>2</sup> | Source |
| --- | --- | --- | --- | --- |
| HumAb | -0.580 | - | 0.336 | Prihoda et al.<sup>2</sup> |
| IgScope fr_mean (geo) | -0.569 | -0.338 | 0.324 | This work |
| IgScope fr_mean (arith) | -0.569 | -0.340 | 0.324 | This work |
| IgScope max_stretch_below_0.5 | +0.568 | +0.360 | 0.322 | This work |
| IgScope chain (arith) | -0.567 | -0.378 | 0.321 | This work |
| HumAb (same-protocol recomputation, H only) | -0.558 | -0.370 | 0.311 | This work (rf-humab 0.0.10) |
| IgScope frac_below_0.5 | +0.558 | +0.371 | 0.311 | This work |
| IgScope fr_mean (min) | -0.557 | -0.334 | 0.310 | This work |
| IgScope residue_mean (arith) | -0.551 | -0.383 | 0.304 | This work |
| IgScope residue_mean (geo) | -0.540 | -0.376 | 0.291 | This work |
| OASis | -0.530 | - | 0.281 | Prihoda et al.<sup>2</sup> |
| ImmunoSeq<sup>11</sup> | -0.530 | -0.409 | 0.281 | Huang et al.<sup>11</sup> |
| IgScope residue_mean (min) | -0.519 | -0.371 | 0.270 | This work |
| IgScope chain (geo) | -0.514 | -0.369 | 0.264 | This work |
| OASis (relaxed)<sup>11</sup> | -0.511 | -0.387 | 0.261 | Huang et al.<sup>11</sup> |
| Germline content | -0.500 | - | 0.250 | Prihoda et al.<sup>2</sup> |
| AntiBERTy fv (arith) | -0.500 | -0.367 | 0.250 | This work |
| AbNatiV<sup>4</sup> | -0.488 | -0.356 | 0.238 | Huang et al.<sup>11</sup> |
| AntiBERTy H only | -0.487 | -0.317 | 0.237 | This work |
| IgScope chain (min) | -0.478 | -0.362 | 0.229 | This work |
| AbLSTM<sup>23</sup> | -0.470 | - | 0.221 | Prihoda et al.<sup>2</sup> |
| MG score | -0.460 | - | 0.212 | Prihoda et al.<sup>2</sup> |
| Z-score | -0.450 | - | 0.203 | Prihoda et al.<sup>2</sup> |
| AntiBERTy fv (geo) | -0.428 | -0.370 | 0.183 | This work |
| T20 | -0.410 | - | 0.168 | Prihoda et al.<sup>2</sup> |
| AntiBERTy L only | -0.391 | -0.329 | 0.153 | This work |
| AntiBERTy fv (min) | -0.390 | -0.372 | 0.152 | This work |
| IgScope cdr_mean (arith) | -0.331 | -0.307 | 0.110 | This work |
| IgScope cdr_mean (geo) | -0.327 | -0.315 | 0.107 | This work |
| IgScope cdr_mean (min) | -0.318 | -0.315 | 0.101 | This work |

**Extended Data Table 3.**
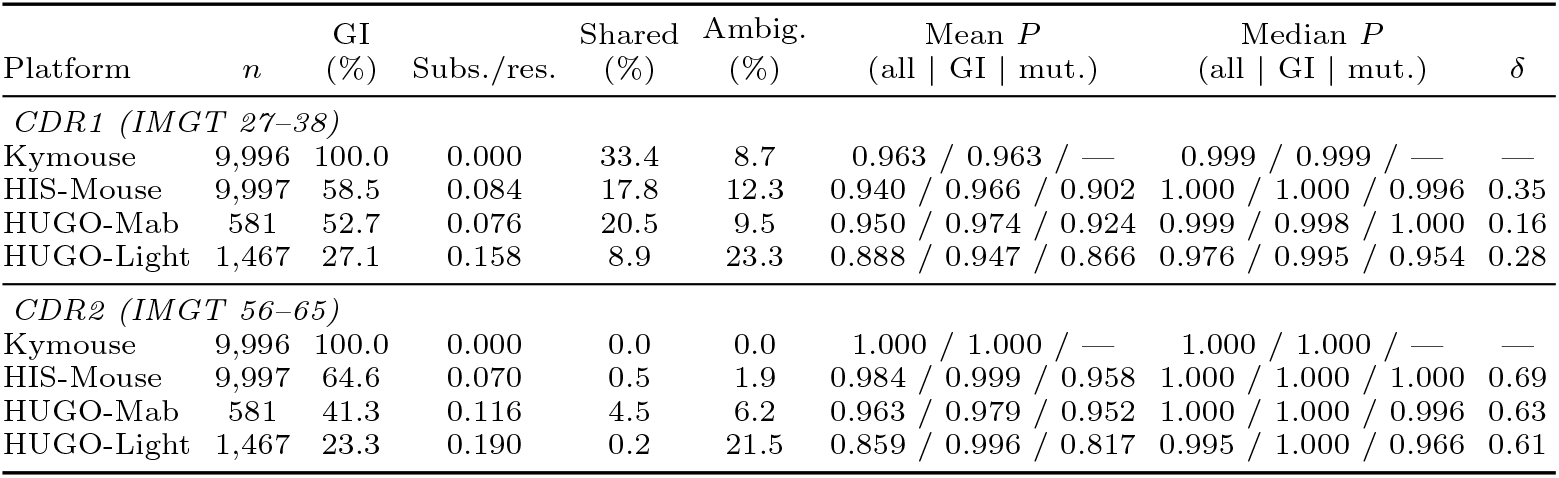
Germline composition and region-score behavior of VH CDR1 and CDR2 across four humanized-mouse platforms. Column headings: **GI (%)**, percentage of chains whose CDR peptide is identical to that of the chain’s own human germline allele; **Subs./res**., substitutions per CDR residue relative to that allele; **Shared (%)**, percentage of peptides identical in at least one human and one mouse germline allele (Supplementary Data 1); **Ambig. (%)**, percentage of chains with 0.5 *≤ P* (human) *≤* 0.9; **Mean *P*** and **Median *P*** (all |GI| mut.), arithmetic mean and median of *P* (human) over all chains in the row and within the germline-identical and mutated strata; ***δ***, Cliff’s delta for *P* (human) of germline-identical versus mutated chains (positive, germline-identical scores higher); **—**, not defined.

## Notes

### Summary of Updates

This revision only fixes formatting errors in the abstract (nonstandard dash characters and minor punctuation/spacing). No scientific content, data, or conclusions have been changed.

