## Supplementary Material for "Multi-resolution humanness scoring of antibodies via a region-isolated language model"

#### Supplementary Information

### 1 Supplementary Tables

Supplementary Table 1 — Composition of the development corpus (train/validation/test).

| Group | Train | Validation | Test | Length (mean) | SHM rate | NP-ins. rate |
|---|---|---|---|---|---|---|
| alpaca-H (camelid) | 6,591,157 | 137,316 | 137,316 | 122.0 | 0.131 | 0.082 |
| camel-H (camelid) | 792,534 | 16,511 | 16,512 | 121.8 | 0.199 | 0.102 |
| human-H | 76,106,133 | 1,585,544 | 1,585,545 | 111.2 | 0.030 | 0.049 |
| human-L (K + L) | 13,746,596 | 286,387 | 286,388 | 108.3 | 0.066 | 0.008 |
| mouse-H | 16,450,962 | 342,728 | 342,729 | 118.6 | 0.010 | 0.027 |
| mouse-L | 529,266 | 11,026 | 11,027 | 108.1 | 0.049 | 0.002 |
| **Total** | **114,216,648** | **2,379,512** | **2,379,517** | | | |

The corpus derives from the Observed Antibody Space (OAS human, mouse and camel repertoires plus 22 alpaca VHH libraries), quality-filtered, de-duplicated within species, reduced by CDR3-grouped Linclust clustering at 95% identity, and split stratified by species x chain type at 95/2/3. human-L pools kappa and lambda chains; alpaca VHHs carry the camelid label. SHM, somatic-hypermutation rate; NP, non-templated insertion rate.

Supplementary Table 2 — Composition of the VDJServer external test set.

| Species | Study | Accession | Files | Chains | H | K | L | Fv length (aa) |
|---|---|---|---|---|---|---|---|---|
| Human | Schmitz et al. (2021) | PRJNA741267 | 30 | 267,492 | 267,492 | 0 | 0 | 121.9 |
| | Benavides et al. (2026) | SUB15840447 | 67 | 34,577 | 34,577 | 0 | 0 | 101.1 |
| Mouse | Valdés-Alemán et al. (2014) | PRJNA275625 | 18 | 137,938 | 137,938 | 0 | 0 | 115.9 |
| | Alameh et al. (2021) | PRJNA669143 | 60 | 64,439 | 64,439 | 0 | 0 | 109.1 |
| | Alsoussi et al. (2020) | PRJNA624801 | 2 | 1,283 | 991 | 286 | 6 | 116.6 |
| Camelid | White et al. (2026), NanoMAP | PRJNA1466709 | 211 | 499,752 | 499,752 | 0 | 0 | 119.7 |
| **Total** | 6 studies | | 388 | **1,005,481** | 1,005,189 | 286 | 6 | 118.4 |

The VDJServer external test set is a BCR (IGH, IGK and IGL) subset of the VDJServer AIRR Data Commons [7, 8], a public repository that contained 39 studies, 3,408 repertoires and approximately 2.5

billion annotated rearrangements at its December 2021 release; TCR records in the same repository were discarded. Chains are IMGT-numbered Fv amino-acid sequences retained after translation of the deposited nucleotide records and filtering for productive, in-frame BCR rearrangements of 100–148 aa; exact duplicates were removed within and across files, sequences were clustered at 95% Fv identity (mmseqs2 linclust) and only cluster centers were IMGT-numbered, retaining chains with CDR1 $\geq$ 5, CDR2 $\geq$ 1 and CDR3 $\geq$ 5 residues. From 23,322,905 deposited sequences, 7,718,055 remained after quality control and deduplication and 4,206,938 clusters were obtained (Human 303,453; Mouse 210,152; Camelid 3,693,333, of which 500,000 cluster centers were sampled); 1,005,481 chains were retained. Two studies present in the repositories were excluded: CDR3-only amplicons shorter than a complete Fv, and FR2-anchored amplicons lacking CDR1. Alpaca libraries contribute VHH heavy chains only. Fv length, mean number of IMGT-numbered residues.

Supplementary Table 3 — ADA $\leq$ 10% versus $>$ 10% binary classification on the deduplicated 210-antibody ADA217 set (positives 48, 22.9%).

| Method | ROC-AUC | AUPRC | F1 | Threshold |
|---|---|---|---|---|
| **IgScope (chain P(human))** | **0.807** [0.730, 0.876] | **0.605** [0.469, 0.746] | **0.652** | MCC-optimal 0.46 |
| MixMHC2pred-2.0 [11] | 0.793 | 0.565 | 0.523 | as reported |
| HLAIIPred [10] | 0.784 | 0.598 | 0.533 | as reported |
| NetMHCIIpan-4.1 [12–13] | 0.767 | 0.487 | 0.514 | as reported |
| Graph-pMHC [14] | 0.760 | 0.558 | 0.493 | as reported |

Bracketed values are 2,000-replicate bootstrap 95% confidence intervals (IgScope only; not reported for the literature methods). Rows are ordered by ROC-AUC. The set follows the cleaning protocol of Haghighatlari et al. [10]: of five duplicate heavy-light pairs, three on the same side of the 10% threshold contribute one representative each and two conflicting pairs are removed entirely (217 $-$ 7 = 210). The IgScope classification threshold was selected on this evaluation set (MCC-optimal); the literature F1 values use thresholds selected by the original authors. ROC-AUC and AUPRC are threshold-independent.

#### 2 Supplementary Figures

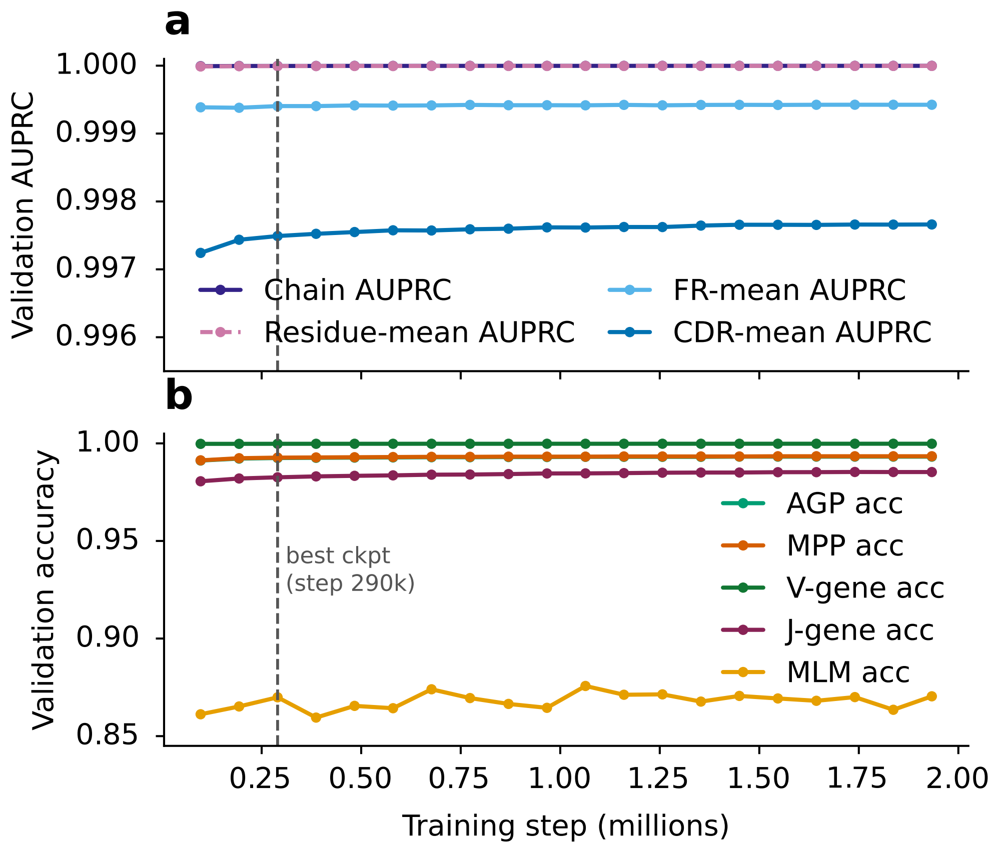


**Figure S1. Training convergence.** a, AUPRC of the chain, CDR-mean, FR-mean and residue heads; b, AGP, MPP, V-gene, J-gene and MLM accuracy. Metrics were evaluated every half epoch on the full validation set against training step; the dashed line marks the best checkpoint (epoch 1, step 290,001) selected by validation CDR-mean AUPRC (0.9975). Chain, FR-mean and residue AUPRC remain above 0.9993 and V-gene accuracy above 0.9997 throughout training.

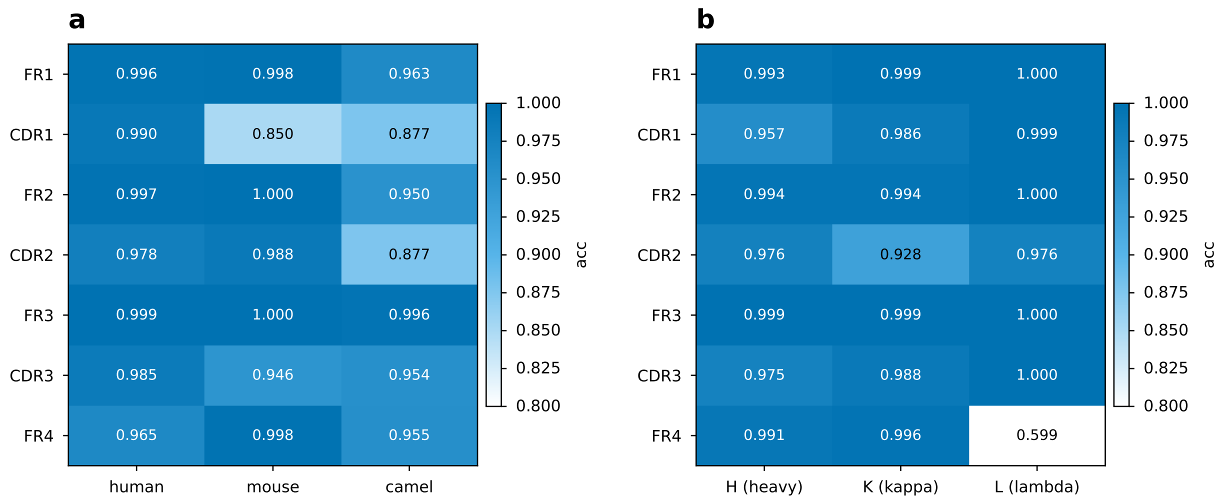


**Figure S2. Region accuracy heatmaps.** a, per-IMGT-region species-classification accuracy by species (human, mouse, camelid). b, per-IMGT-region accuracy by chain type (H heavy, K kappa, L lambda). Test set n = 2,379,517 natural chains.

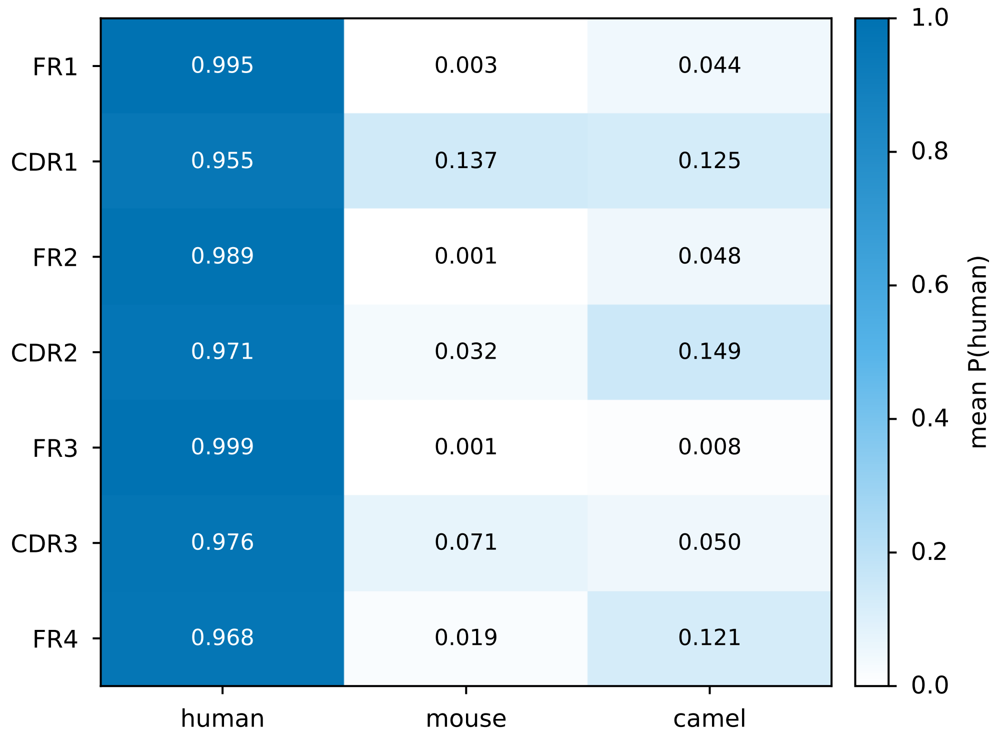


**Figure S3. Example multi-resolution humanness profiles.** Mean chain-level P(human) per IMGT region and species; human frameworks score near 1.0, whereas mouse and camelid frameworks score near zero, with the highest camelid values in CDR3.

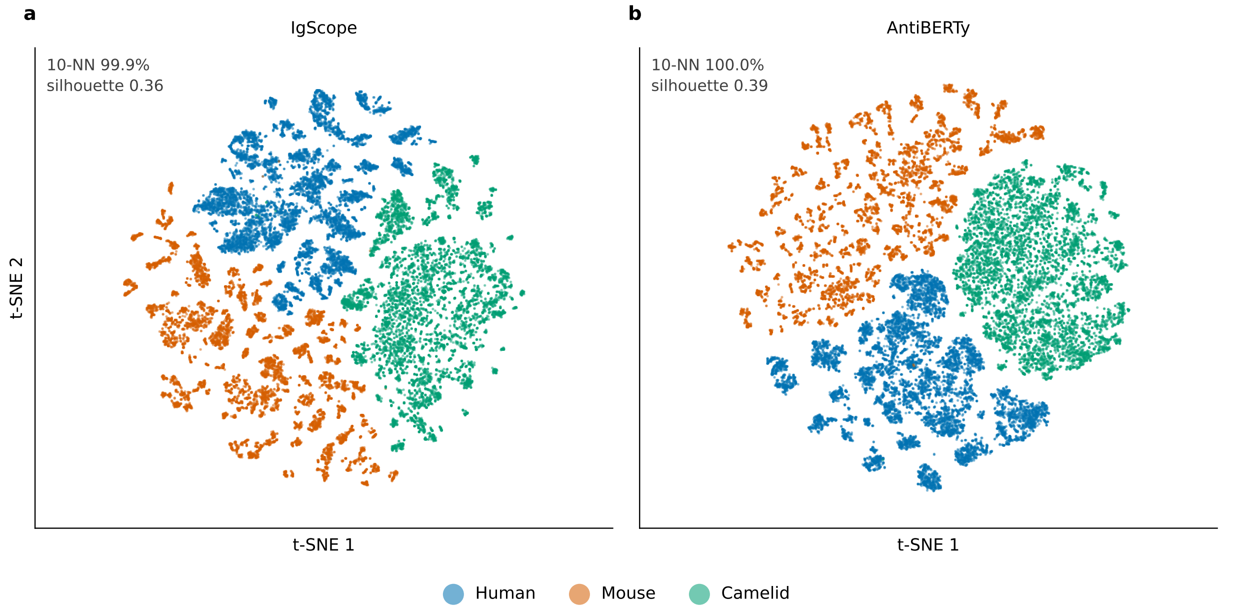


**Figure S4. t-SNE map of whole-chain embeddings by species. a**, IgScope; **b**, AntiBERTy. A 30,000-chain species-stratified subsample of the external test set (10,000 chains per species) was encoded as whole-chain [CLS] embeddings (480 dimensions for IgScope, 512 for AntiBERTy) and projected to two dimensions with t-SNE (50 principal components, perplexity 30, seed 42; identical settings for both models). Points are colored by species (Okabe–Ito palette). Values in the upper left of each panel give the fraction of chains whose 10 nearest neighbors share their species label and the silhouette coefficient. Both models place the three species in separate regions of the map.

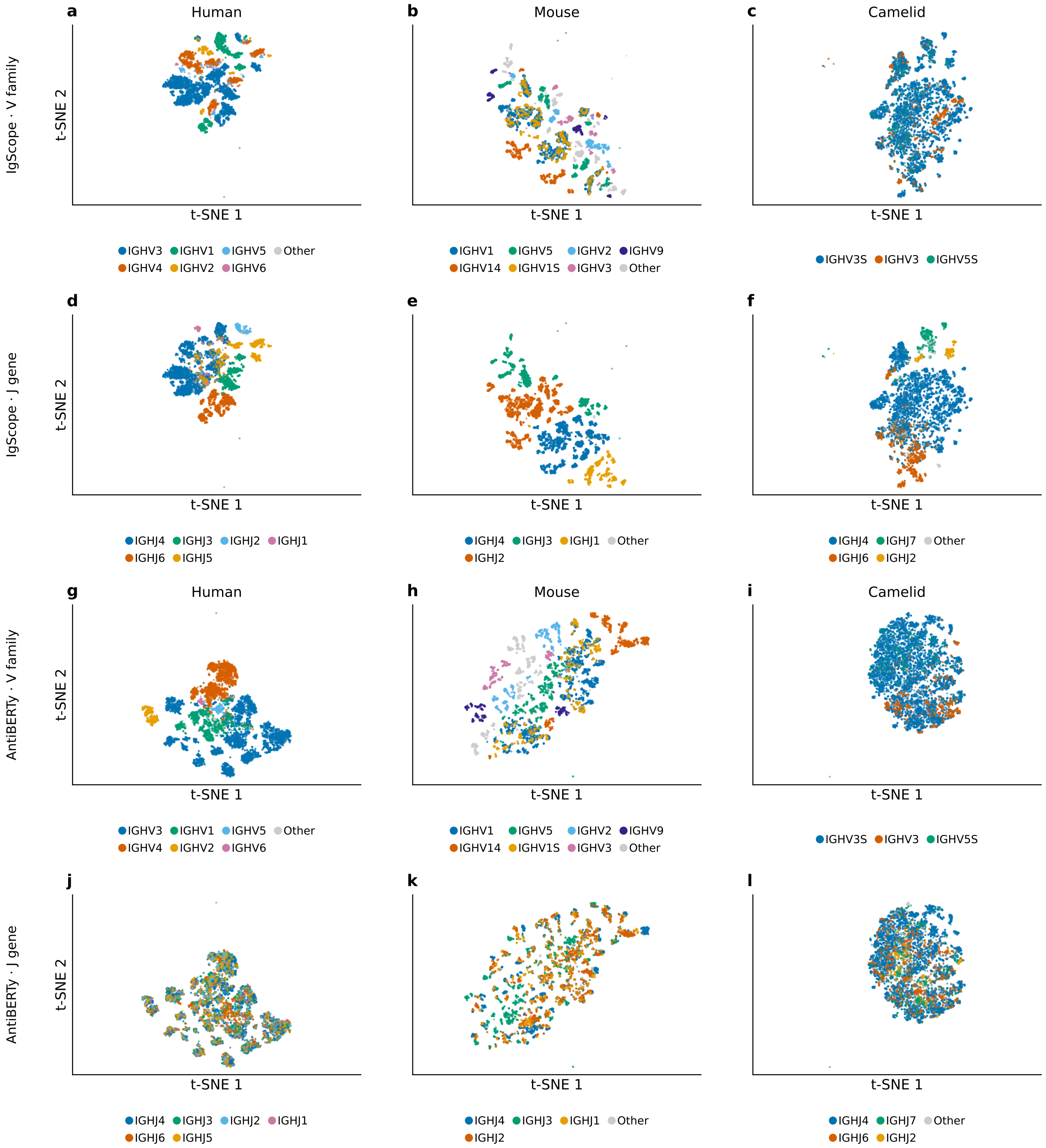


**Figure S5. V-family and J-gene stratification of the t-SNE islands.** The embeddings of Fig. S4 colored by assigned V family (rows 1 and 3) or J gene (rows 2 and 4) for IgScope (**a–f**) and AntiBERTy (**g–l**); columns are human, mouse and camelid. Families carried by fewer than 50 (human), 500 (mouse) or 100 (camelid) chains are pooled as Other (gray). Within-species 25-nearest-neighbor label purity exceeds the chance level ($\sum p^2$) in every panel; for J genes it is 0.929/0.997/0.914 for IgScope versus 0.544/0.700/0.769 for AntiBERTy (human/mouse/camelid; chance 0.327/0.290/0.587), so IgScope's finer sub-islands follow V–J usage whereas AntiBERTy's largely do not.

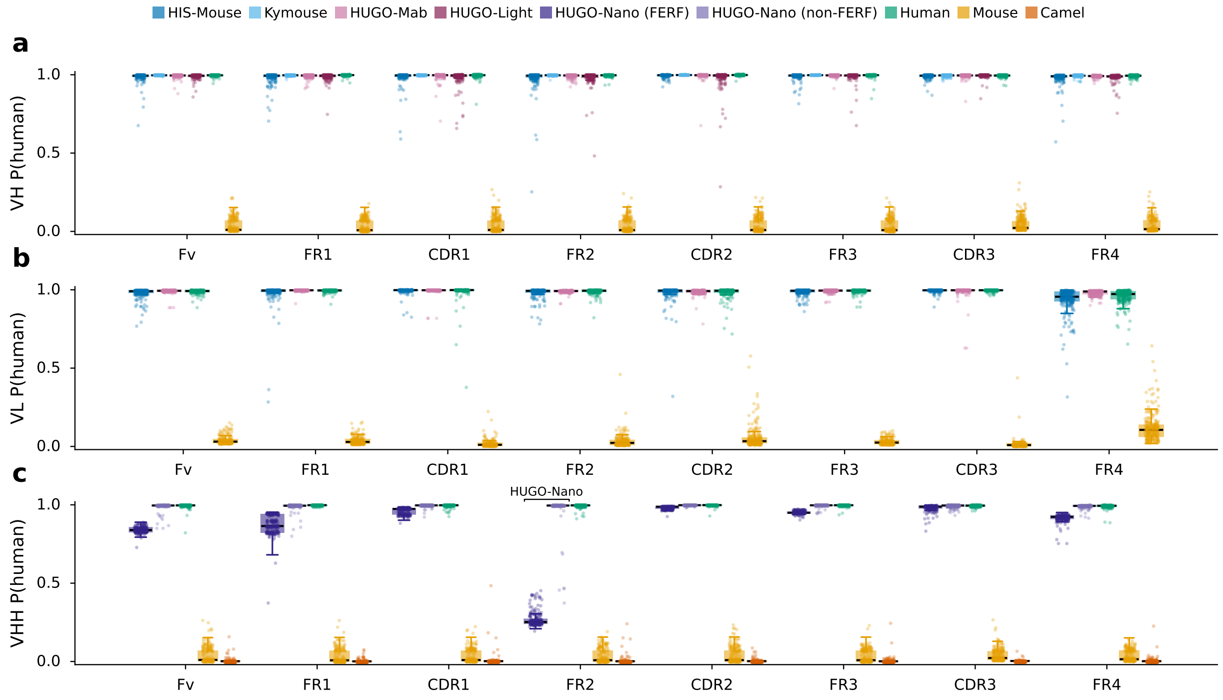


**Figure S6. Region scores from the region head versus the mean of residue-level scores, for humanized-mouse platforms.** Same layout as Fig. 6a–c, with HUGO-Nano chains split as in Fig. 6c; region values are the arithmetic mean of per-residue P(human) within each region. For the FERF subset of HUGO-Nano the two calibres differ markedly (region-head median 0.001; residue-mean median 0.252): the region score is dominated by the four engineered camelid hallmark residues, while residue scores at the remaining FR2 positions — depressed beyond the engineered sites, consistent with context effects propagated from those positions — hold the residue mean well above zero.

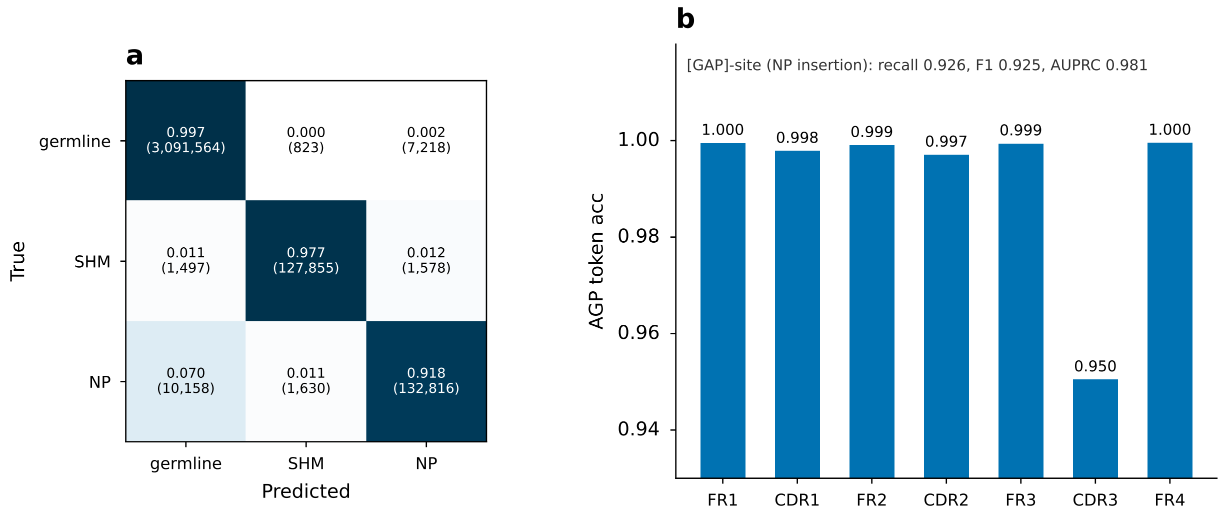


**Figure S7. AGP/MPP diagnostics.** a, MPP three-class (germline / SHM / NP) confusion matrix on a 30,000-chain species-stratified subsample of the test set (rows normalized; counts in parentheses; subsample MPP accuracy 0.9932 versus 0.9927 on the full test set). b, AGP germline-reconstruction token accuracy per IMGT region; the [GAP]-site (NP insertion) detection scores are from the full test set (recall 0.926, F1 0.925, AUPRC 0.981). Per-class MPP F1 values are reported in Extended Data Table 1.

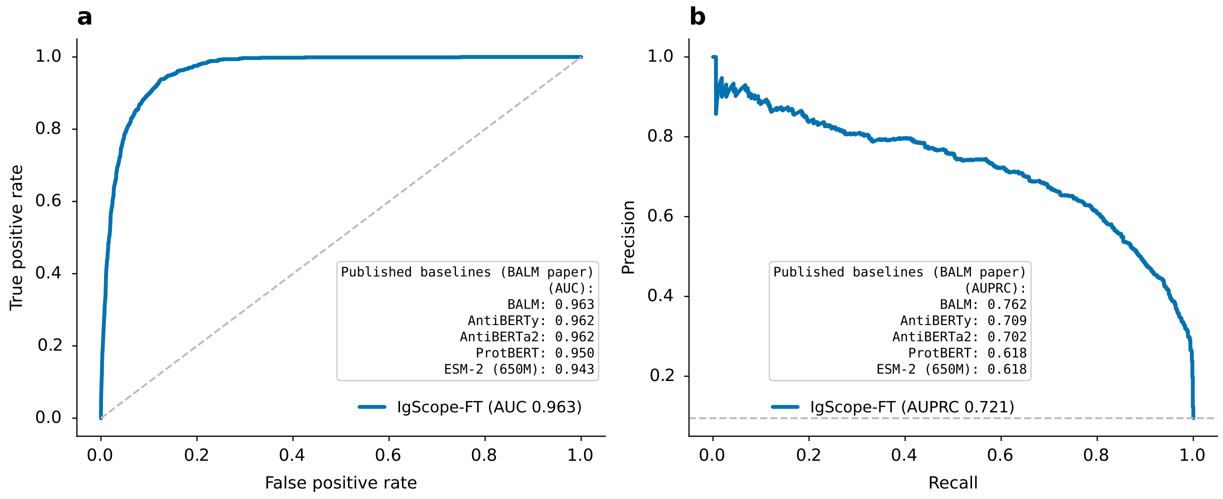


**Figure S8. Paratope prediction on SAbDab (BALM split).** a, receiver operating characteristic; b, precision–recall (AUPRC). IgScope-FT-v1 per-residue scores (F1 0.689 at the validation-optimal threshold 0.85; AUC 0.963); published baseline values are from the BALM paper under the same split.

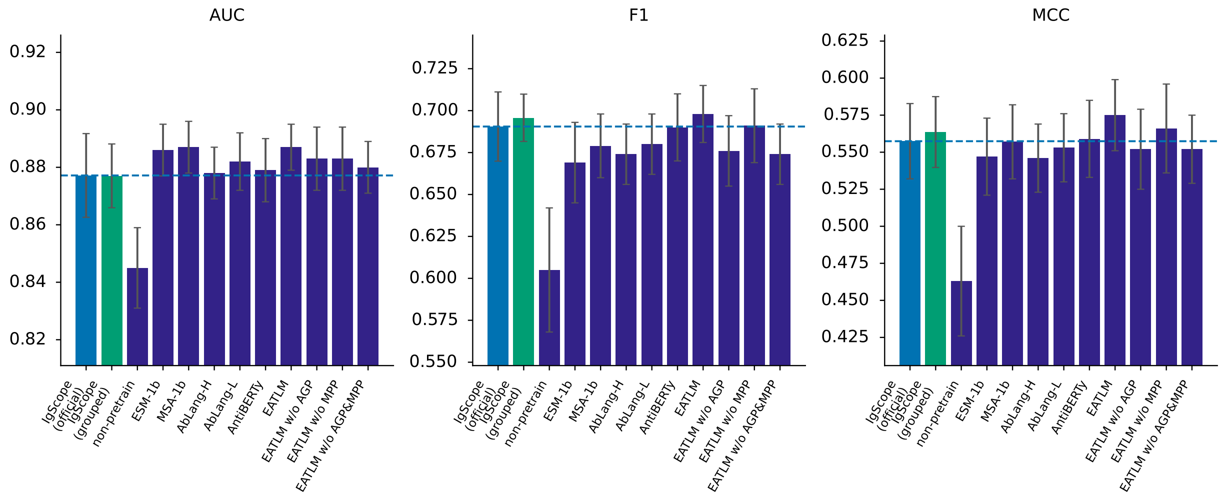


**Figure S9. Paratope prediction on ATUE** (10-fold sequence-grouped; F1 0.696, statistically tied with best baseline EATLM 0.698).

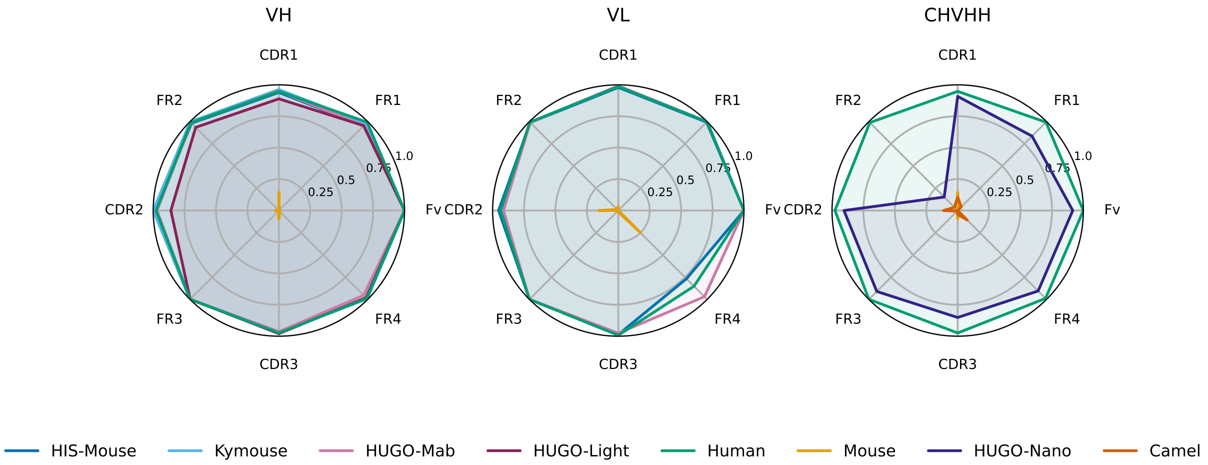


**Figure S10. Humanized-mouse strain radar summary** (four strains humanized throughout; sole gap: HUGO-Nano chVHH FR2).

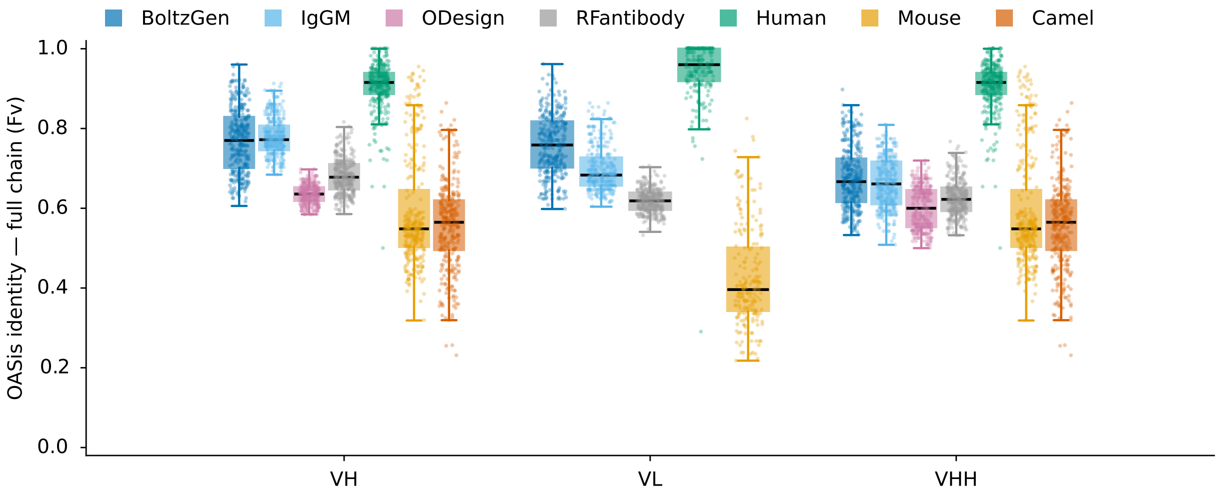


**Figure S11. OASis Fv scores on the same de novo designs** (under-separates designs from natural controls, $\rho$ = 0.561 versus IgScope region scores).
